# Single-cell chromatin and gene-regulatory dynamics of mouse nephron progenitors

**DOI:** 10.1101/2021.10.13.464099

**Authors:** Sylvia Hilliard, Giovane Tortelote, Hongbing Liu, Chao-Hui Chen, Samir S. El-Dahr

## Abstract

**Background:** Cis-regulatory elements (CREs), such as enhancers and promoters, and their cognate transcription factors play a central role in cell fate specification. Bulk analysis of CREs has provided insights into gene regulation in nephron progenitor cells (NPCs). However, the cellular resolution required to unravel the dynamic changes in regulatory elements associated with cell fate choices remains to be defined.

**Methods:** We integrated single-cell chromatin accessibility (scATAC-seq) and gene expression (scRNA-seq) in embryonic E16.5 (self-renewing) and postnatal P2 (primed) mouse Six2^GFP^ NPCs. This analysis revealed NPC diversity and identified candidate CREs. To validate these findings and gain additional insights into more differentiated cell types, we performed a multiome analysis of E16.5 and P2 kidneys.

**Results:** CRE accessibility recovered the diverse states of NPCs and precursors of differentiated cells. Single-cell types such as podocytes, proximal and distal precursors are marked by differentially accessible CREs. Domains of regulatory chromatin as defined by rich CRE-gene associations identified NPC fate-determining transcription factors (TF). Likewise, key TF expression correlates well with its regulon activity. Young NPCs exhibited enrichment in accessible motifs for bHLH, homeobox, and Forkhead TFs, while older NPCs were enriched in AP-1, HNF1, and HNF4 motif activity. A subset of Forkhead factors exhibiting high chromatin activity in podocyte precursors.

**Conclusion:** Defining the regulatory landscape of nephrogenesis at single-cell resolution informs the basic mechanisms of nephrogenesis and provides a foundation for future studies in disease states characterized by abnormal nephrogenesis.

**Significance Statement:**

- Nephron progenitor cells (NPCs) are a multipotent population giving rise to all cell types of the nephron. At any given time, the NPC’s choice to self-renew or differentiate is determined not only by its transcription factor (TF) repertoire but also by the genome accessibility of the cognate cis-regulatory elements.
- Using single-cell analysis, we demonstrate the heterogeneity of NPCs at the epigenetic level and observe dynamic and cell type-specific changes in chromatin accessibility. Fate-determining TFs harbor domains rich in interactive chromatin that are established prior to gene activation.
- These findings illustrate the importance of chromatin-based mechanisms in the regulation of nephrogenesis and may have implications for nephron regeneration and repair.

## INTRODUCTION

In the developing kidney, nephron progenitor cells (NPCs) reside in a niche called the cap mesenchyme, a crescent-shaped cellular compartment surrounding the adjacent ureteric branch tip. Based on molecular marker analysis, the cap mesenchyme is divided into a self-renewing Cited1^+^/Six2^+^ compartment, a transit Cited1^-^/Six2^+^ compartment, and a late Six2^low^/Wnt4^+^ compartment called the pretubular aggregate, the precursor of the renal vesicle, and the earliest epithelial precursor of the nephron (1, 2). Single-cell transcriptomics of human and mouse kidneys refined the signatures of NPC states and delineated the developmental trajectories and the potential transcriptional regulatory networks associated with cell fate decisions (3–7). The NPC’s choice to self-renew or differentiate and gain a new identity is determined not only by its transcription factor (TF) repertoire but also by the accessibility of the cognate cis-regulatory elements (CREs). Accordingly, to fully characterize the NPC states and fates, it is necessary to have an integrated view of chromatin and gene expression states across cell types, developmental trajectory, and life span.

Chromatin accessibility is a dynamic process that drives tissue development by permitting or restricting access of TFs to cognate CREs. ATAC-seq (<u>A</u>ssay for <u>T</u>ransposase <u>A</u>ccessible <u>C</u>hromatin) has become the assay of choice to decipher open (accessible) chromatin domains composed of promoters and enhancers (8). We previously reported using bulk ATAC-seq in native embryonic and neonatal mouse Six2^GFP^ NPCs that NPCs undergo age-dependent changes in chromatin accessibility at active enhancers of renewal and differentiation genes (9). These differential dynamic chromatin states were also observed in expanding NPCs ex vivo, suggesting that young and old NPCs are intrinsically different at the epigenetic level (9). Changes in chromatin accessibility to TCF/LEF factors at pro-differentiation enhancers were also shown to mediate gene expression in response to ß-catenin activation (10), underscoring the importance of concerted actions of chromatin and transcription factors in gene regulation. However, given the dynamic changes in NPC states that occur during differentiation, it is difficult to accurately assign a chromatin state to a specific cell population using bulk analysis. In a recent study to address this issue (11) the investigators performed snRNA/snATAC-seq in newborn and adult mouse kidneys revealing key developmental regulators of NPC commitment decisions to the podocyte, proximal and distal fates and linking single nucleotide variants associated with human kidney disease to regulatory elements in key developmental genes. In the present study, we integrated single-cell chromatin accessibility and gene expression profiles obtained from embryonic and neonatal mouse NPCs and subsequently validated our singleomes using single-cell profiles derived from the same cell (multiome). These paired maps recovered a class of genes, frequently lineage-determining transcription factors, enriched in putative enhancers whose accessibility was strongly linked to gene expression.

## MATERIALS AND METHODS

Animal protocols utilized in this study were approved by and in strict adherence to guidelines established by the Tulane University Institutional Animal Care and Use Committee.

### Mice

Six2^GFP^ NPCs were isolated from kidneys of *embryonic day* E16.5 (n=3) and *postnatal day* P2 (n=4) *Six2GFPCre* (Six2^TGC^) mice bred on a mixed genetic background by fluorescent-activated cell sorting (FACS), as described (9). For multiomics, cells were isolated from kidneys of E16.5 and P2 C57Bl/6 mice.

### Single-cell ATAC seq data generation from E16.5 and P2 Six2^GFP^ kidneys

Nuclei were prepared from freshly explanted kidneys in accordance to the protocol outlined in CG000169 | RevB (10X Genomics). Approximately 100,000 dissociated cells were washed in ice-cold 1X PBS supplemented with 0.04% BSA. Cells were spun down at 500g for 5min at 4°C. The cell pellet was re-suspended by gentle pipetting in 45ul Lysis buffer 10 mM Tris pH 7.4, 10 mM NaCl, 3 mM MgCl2 0.1% NP-40, 0.1% Tween-20, 0.01% Digitonin and 1% BSA. Lysis was allowed to proceed on ice for 5 minutes after which 50ul of ice-cold wash buffer (10 mM Tris pH 7.4, 10 mM NaCl, 3 mM MgCl2, 0.1% Tween-20 and 1% BSA) was added before spinning down again and re-suspending in 45 μL 1X Nuclei Dilution Buffer (10x Genomics). After the last centrifugation step, 7ul of 1X diluted Nuclei Buffer was added to the nuclei pellet. Two microliters of the nuclei preparation plus 8ul of diluted Nuclei buffer was stained with 10ul of AO/PI staining solution (Nexcelom, MA). Both concentration and viability of the sample was read using the Cellometer Auto 2000 Cell Viability Counter (Nexcelom, MA). The remaining 5ul of nuclei was adjusted for a targeted nuclei recovery of 5000 before proceeding to the Chromium Single Cell ATAC Reagents and Gel Bead Kit protocol CG000168 (PN1000111,10x Genomics).

Paired-end dual index libraries were sequenced on the Illumina NextSeq 550 sequencer. Raw data was extracted and processed using the Cell Ranger pipeline (v1.2.0, 10X Genomics) and the mm10 Mus musculus reference assembly.

### Multiome data generation

Cells were isolated from mouse kidneys explanted at E16.5 and P2. The kidneys were dissociated by incubation in Accutase (AT104, Innovative Cell Technologies, Inc.) with slow rotation at 37°C for up to an hour with intermittent gentle pipetting. The reaction was stopped with FBS and the cells were spun down at 500rpm for 5min. The cell pellet was resuspended in 1XPBS supplemented with 10mM EDTA and 2%FBS. Following resuspension, the cells were passed through a 40um cell strainer to obtain a single cell suspension (BD Falcon, catalog# 352235).

Nuclei isolation was achieved with 800,000 cells per age group using the protocol CG000366 RevB (10XGenomics) established for mouse tissues. To achieve cell lysis, we incubated the cells in 0.1X Lysis Buffer for 8min on ice. After the wash and centrifugation steps, the nuclei pellet was resuspended in cold 1X Nuclei Buffer. Approximately 6000 nuclei per sample were used to prepare the libraries in accordance with the steps detailed in the Chromium Next GEM Single Cell Multiome ATAC + Gene Expression protocol (CG000338, RevD) and the 10X Reagent Kit PN-1000284. Briefly, nuclei suspensions were incubated in a Transposition Mix to transpose and barcode accessible DNA fragments. The transposed nuclei were partitioned into nanoliter-scale Gel Bead-In EMulsion (GEMs). The barcoded transposed DNA and barcoded full-length cDNA from poly-adenylated mRNA were generated and amplified by PCR to obtain sufficient complexity for both ATAC and gene expression library constructions. P7 and a sample index were added to transposed DNA during ATAC library construction via PCR. Barcoded cDNA enzymatic fragmentation, end-repair, A-tailing, and adaptor ligation were followed by PCR amplification. Both the ATAC and 3′-Gene Expression libraries generated contained standard Illumina P5 and P7 paired-end constructs. Library quality controls were performed with an Agilent High Sensitivity DNA kit on the Agilent 2100 Bioanalyzer together with quantitation on a Qubit 2.0 fluorometer. Libraries were sequenced separately with individual parameter settings. Pooled libraries at a final concentration of 650pM were sequenced with paired end dual index configuration on an Illumina NextSeq 550 using Illumina High-Output 150 cycle kits (Cat No.20024907). Cell Ranger-arc version 1.0.1 software (10X Genomics) was used to process raw sequencing data for further downstream analyses.

### Analysis using ArchR package

Fragment files generated with cell-ranger -arc version 1.0.1 pipeline (10X Genomics) were used as input for ArchR (Analysis of Regulatory Chromatin in R) (archproject.com)(12). The input fragment file was processed as chunks per chromosome and stored in HDF5 format (hierarchical data format version 5) allowing rapid access and efficient read and write functions during analysis. The QC steps involved the removal of all low-quality cells and predicted heterotypic doublets from our analysis. All cells with a TSS enrichment score greater than or equal to 4 and with ATAC fragment counts greater than 1000 but less than 100,000 per cell were retained in our analysis.

A chromatin accessibility matrix was generated using a genome-wide, non-overlapping, tile matrix of 500bp bins. The gene score algorithm allowed the prediction of gene activity/expression based on ATAC-seq data. It measures accessibility at promoter plus gene body and distal regulatory elements at weighted distances from the gene TSS and TTS while accounting for gene size differences and neighboring gene boundaries. We used a default of 100kb window flanking the gene.

To obtain a low dimensional representation of single-cell ATAC datasets in terms of principal components and UMAP coordinates, we applied an iterative latent semantic indexing approach (13, 14). In brief, Dimensionality reduction was achieved by TF-IDF normalization and Singular Value Decomposition (SVD)/Latent Semantic Indexing (LSI) through user-defined iterations. The first iteration uses top accessible features while the second uses top variable features. Clustering is achieved by Shared Nearest Neighbor algorithm implemented by Seurat followed by 2D embedding in the UMAP (Uniform Manifold Approximation and Projection) space for visualization. Peak analysis was preceded by calling pseudo-bulk replicates of each cluster using default parameters. MACS2 was used for peak calling and the generation of a non-overlapping union peak matrix. ChromVAR (15) (v.1.6) was used to obtain TF accessibility profiles using position weight matrices from the CisBP mouse_pwms_v2 database (16).

Co-accessibility detection in ArchR follows an approach introduced by Cicero (17) by creating low-overlapping aggregates of single cells. ArchR uses optimized methods for calculating significantly correlated peak-peak as well as cis-regulatory peak-to-gene associations (correlation cutoff = 0.45, resolution =1000). The primary difference between peak-to-gene links and peak-peak co-accessibility is that co-accessibility correlates accessibility between two peaks (ATAC-seq only analysis), while peak-to-gene linkage leverages integrates scRNA-seq data to correlate peak accessibility and gene expression which may be a more pertinent approach to identifying gene regulatory interactions.

We applied the GetAssayData function (Seuart 4.0 Package) to recover the log-normalized information stored at the data slot of the RNA assay from the E16.5 Seurat object. This information was used to construct the expression matrix used for regulatory network analysis. Next, to identify TFs and characterize cell states, we employed cis-regulatory analysis using the R package SCENIC v1.1.1 (Aibar et al., 2017), which infers the gene regulatory network based on co-expression and DNA motif analysis. The network activity is then analyzed in each cell to identify recurrent cellular states. We used GENIE3 to identify TFs were identified and compiled into modules (regulons), which were subsequently subjected to cis-regulatory motif analysis using RcisTarget mm9 files with two gene-motif rankings: 10 kb around the TSS and 500 bp upstream. Regulon activity in every cell was then scored using AUCell. Finally, binarized regulon activity was projected onto Seurat-create UMAP clustering.

### RNAscope in situ hybridization

RNAscope was applied to formalin-fixed, paraffin-embedded tissue sections from E16.5 kidneys using the Multiplex Fluorescent Assay (Advanced Cell Diagnostics, Cat# 323100) following the manufacturer’s recommendations. The list of probes used are as follows: Foxl1 (Cat# 407401), Foxp1 ((Cat#485221), Foxn3 (Cat# 586011), Nabp1 (Cat#1049221), Mcmdc2 (Cat#477971).

## RESULTS AND DISCUSSION

### CRE accessibility reveals NPC precursor states and novel cellular markers

To elucidate the cellular heterogeneity of nephron progenitors, we created gene-regulatory datasets using the Chromium platform (10x Genomics) to generate single-cell ATAC-seq (scATAC) and single-cell RNA-seq (scRNA) libraries from fluorescence-activated sorted Six2^GFP^ cells freshly harvested from E16.5 and P2 kidneys of transgenic Six2-GFP-Cre (Six2^TGC^) mice. We reasoned that while embryonic NPCs are engaged in self-renewal, neonatal NPCs are primed for differentiation nearing the time of cessation of nephrogenesis (P3-P4 in mice). Overall, we obtained 13,506 (E16.5) and 7,232 (P2) single-cell epigenomes and 10,544 (E16.5) and 7,250 (P2) single-cell transcriptomes after quality control and filtering (**Figure S1, A,B and Figure S2 A,B**). After data processing using the Cell Ranger pipeline (10x Genomics), we employed ArchR (12) for downstream pipeline analysis, as described in the Methods section.

To reveal global similarities and differences between individual cells, we performed dimension reduction using uniform manifold approximation and projection (UMAP) and clustering. We employed the latent semantic indexing (LSI) approach for scATAC to obtain a low-dimensional embedding, cell clustering and consensus sets of 753614 (E16.5) and 309584 accessible peaks representing potential CREs. We next called clusters in both scATAC and scRNA (GSE124804) datasets and annotated these clusters using gene activity scores (a metric defined by the aggregate local chromatin accessibility of genes) [**Methods**] and gene expression (**Figure 1 A, B, E, F**). Integrating the derived gene activity scores with gene expression levels using canonical correlation analyses (CCA) to match cells from one modality to their nearest neighbors in the other (**Figure 1 C, G**) showed that the major NPC states were recovered at both E16.5 and P2. Thus, chromatin accessibility is comparable to gene expression in predicting the major cell states of NPCs.

**Figure 1.**
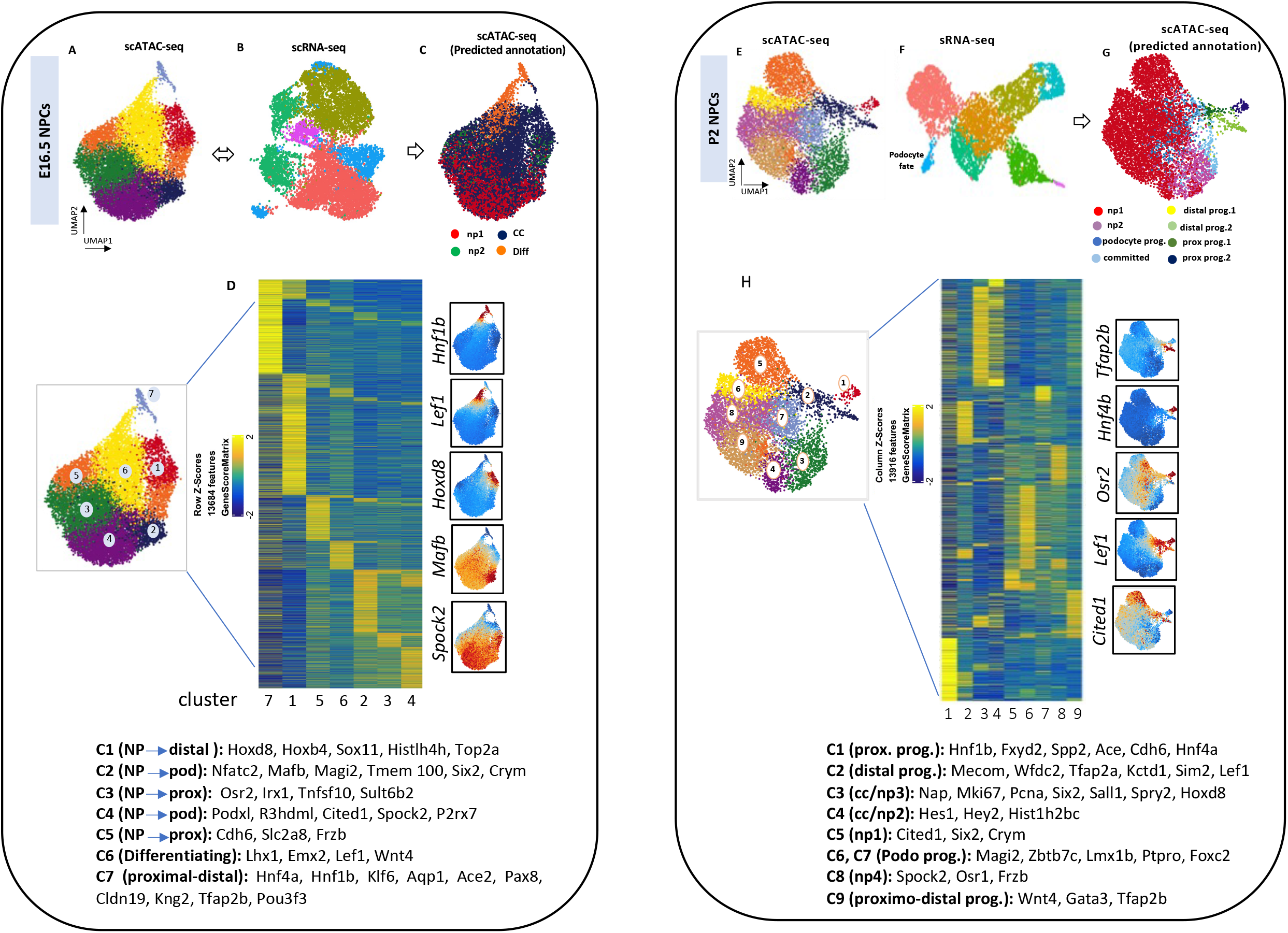
Integrated single-cell ATAC/RNA-seq reveals diversity of cell states in E16.5 and P2 Six2^GFP^ NPCs. (A,E) UMAP plots of scATAC datasets with chromatin accessibility (gene scores)-based cell-type assignments in E16.5 (A) and P2 (E) NPCs. (B,F) UMAP plots of scRNA-based cell-type assignments. (C,G) UMAP plots of integrated scATAC/scRNA-based cell clustering. (D,H) Heatmaps of scATAC gene scores for the top 13684 (E16.5) and 13916 (P2) accessible peaks. Representative genes of each cluster are shown. NP: nephron progenitor; CC cell cycling; Pod: podocyte

We examined the gene scores matrix (derived from chromatin accessibility) to determine whether we can identify precursor cell states in the embryonic stage NPC. The results revealed that chromatin accessibility can identify precursor states of podocyte, proximal and distal fates (**Figure 1 D**). For example, Cluster 1, which contains markers of cell cycle genes, also features the distal tubule fate marker Hoxd8, whereas clusters 3 and 5 feature markers of proximal progenitors Osr2 and Cdh6. The Clusters 2 and 4, representing early nephron progenitors, feature markers of podocytes. Similarly, progenitor cell states were also identified at P2 based on chromatin accessibility gene scores (**Figure 1 H**), although by this advanced stage of maturation scATAC-seq and scRNA-seq were equivalent in identifying precursor states. Collectively, these findings demonstrate that single-cell chromatin accessibility offers a sensitive means to recover precursor cell states in fetal NPCs.

scATAC-seq also allowed the detection of novel markers, Mcmdc2 (Minichromosone Maintenance Domain Containing-2) and Nabp1(Nucleic acid-binding protein-1), showing selective chromatin accessibility in proximal or distal cell precursors, respectively (**Figure S3 A,C**). Using RNAScope in situ hybridization, we confirmed the RNA expression of *Mcmdc2* in proximal (LTL-positive) tubules, whereas *Nabp1* was expressed in distal (LTL-negative) segments (**Figure S3 B, D**). Interestingly, both genes are involved in DNA repair pathways.

We used the R package Cicero (17) to predict cis-regulatory chromatin interactions for individual cell types. Cis co-accessibility networks (CCAN) are families of chromatin regions that co-vary in their accessibility and can be used to predict chromatin interactions. Within NPCs at E16.5 and P2, most Cicero connections were either within a promoter region or between a promoter and another location like introns and intergenic regions (**Figure S4 A-C**). Overall, there were twice as many CCANs in P2 than E16.5 NPCs reflecting increasing complexity of transcriptional regulation with maturation.

### CRE-Gene associations identify potential regulatory chromatin

Because co-variations in chromatin co-accessibility generally have only modest correlations with gene expression, we linked distal peaks to genes in *cis*, based on co-variation in chromatin accessibility and gene expression across all cells to identify CRE-gene pairs during NPC differentiation. Ma and colleagues (18) called chromatin domains enriched in more than 10 CRE-gene associations <u>D</u>omains <u>o</u>f <u>R</u>egulatory <u>C</u>hromatin (DORCs). DORCs identified lineage-determining genes in the hair follicle that are accessible prior to onset of gene expression. Trevino and colleagues (BioxRv 2021, pre-print) referred to genes enriched in more than 10 peak-gene links as <u>G</u>enes with <u>P</u>redictive <u>C</u>hromatin (GPCs); GPCs identified lineage-determining factors during neural corticogenesis. In the present study, we applied similar principles by generating pseudobulk aggregates of matched ATAC/RNA annotations across NPCs linking gene-distal CRE accessibility to gene expression. We identified 7039 and 21788 CRE-gene pairs that represent potential regulatory interactions in E16.5 and P2 Six2^GFP^ datasets, respectively (**Figure 2 A, B**). CRE-gene links correlated strongly with cell type-specific expression of TFs (**Figure S5**). Annotation using the Genomic Regions Enrichment Annotations Tool (GREAT-R) (19) recovered biological processes relating to basic aspects of nephrogenesis ranging from chromatin organization to renal vesicle formation and cell-cell interactions. (**Figure 2 A, B**).

**Figure 2.**
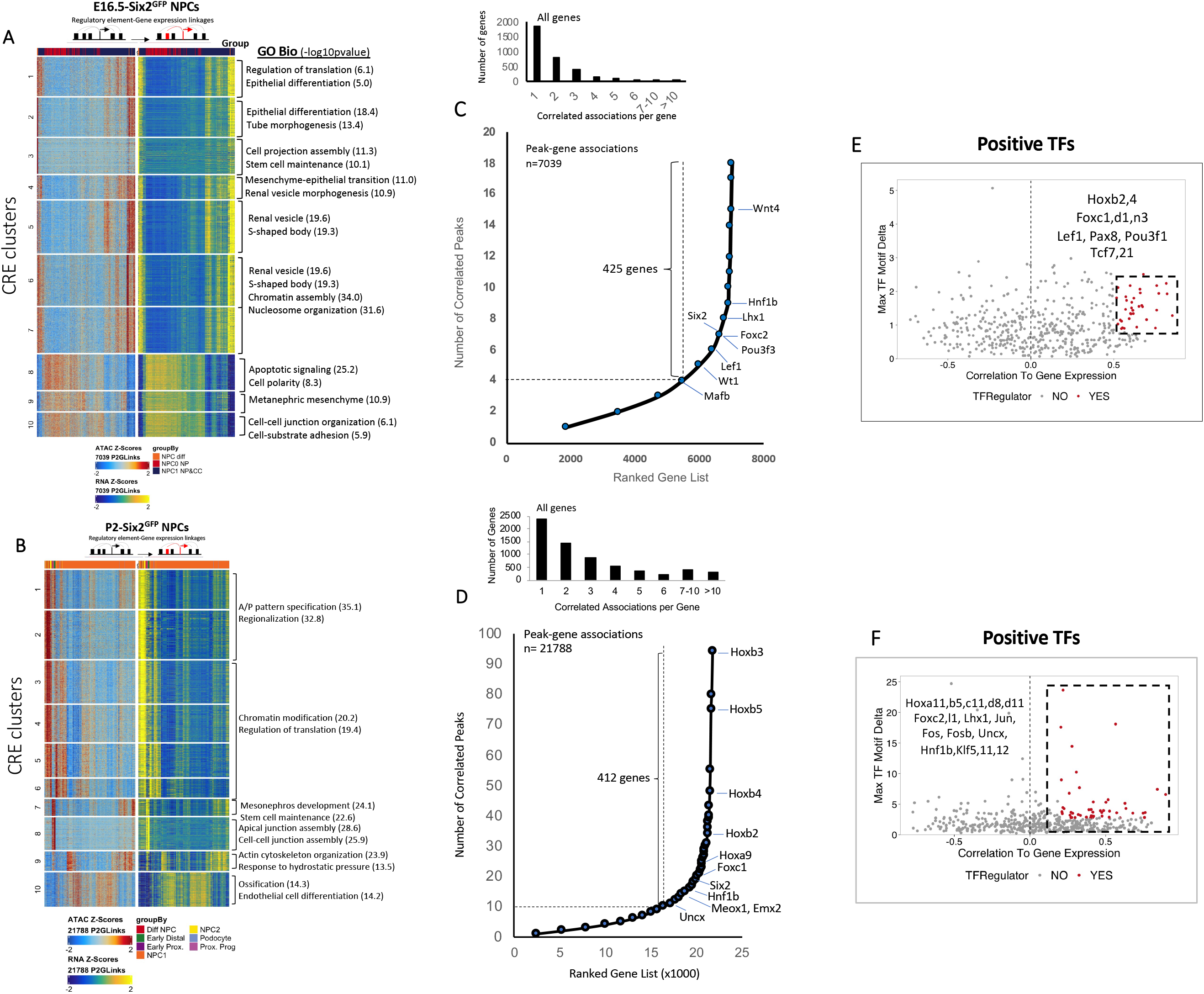
Inferring regulatory chromatin from peak-gene links in Six2^GFP^ NPCs. (A,B) Correlations of 7039 and 21788 peak-gene associations ATAC and RNA Z-scores across all cell states (k means=10). rGREAT annotation of cluster-related biological processes. (C,D) Ranking of genes harboring chromatin domains with a minimum of 4 (E16.5) or 10 (P2) peak-gene links. (E,F) Identification of top positive TF regulators whose correlation between motif and integrated gene expression is greater than 0.5 and 0.2, respectively with an adjusted p-value less than 0.01 and a maximum inter-cluster difference in deviation z-score delta value that is in the top 0.75 quartile.

To determine whether CRE-gene associations have a predictive value in identifying key developmental regulators, we ranked genes across all NPCs based on the number of peak-gene associations and exceeding an inflection point (“elbow”) (18). This analysis yielded 425 (E16.5) and 412 (P2) genes having more than 4 (E16.5) or 10 CRE-gene associations (P2) (**Figure 2 C, D**). These enriched domains of interactive chromatin identified lineage- and fate-determining genes such as *Wnt4*, *Hox*, *Hnf1, Six2, Emx2, Foxc1, Pou3f3* and *Uncx*, among others (**Figure 2 C, D**). We next wished to determine whether genes enriched in interactive domains are concordantly rich in the active enhancer mark H3K27Ac. For this purpose, we leveraged our ChIP-seq data performed on E13.5 and P0 NPCs (9). Notably, 15-25% of genes have interactive chromatin domains overlapping with the top-ranking H3K27Ac-marked enhancers (**Fig. S6**). Of interest, a similar analysis performed in the hair follicle revealed that 34% of genes harboring domains of regulatory chromatin overlapping with H3K27Ac-superenhancers (18), underscoring that active enhancer formation encompasses additional mechanisms.

We next wished to identify “positive” transcriptional regulators whose gene expression scores are highly correlated to their ChromVAR TF motif deviation z-scores in the top quartile (r>0.5, p<0.01). This analysis yielded key lineage-determining genes concerned with NPC identity and fate decisions (**Figure 2 E,F**). Notably, we observed that in differentiation genes, e.g., *Hnf1b,* chromatin accessibility at CREs preceded gene expression (**Figure S7 A**). Indeed, chromatin profiles show accessibility of the *Hnf1b* locus in precursor cells (**Figure S7 B**) indicating the necessity for additional gain of accessible chromatin at the gene’s regulatory elements for cell-type specific expression.

### Defining trajectory of transcriptional regulators in NPCs

To identify TFs that may control the gene expression programs across pseudotime, we linked TF motifs enriched in the different cell states with TF gene expression starting from naïve NPCs and ending in the proximo-distal cluster. At E16.5, Six2, Hoxa9, Hoxc8, and Forkhead box (Foxn3, Foxp1, Foxo1, Foxc1, Foxd1) are the initial TFs to engage chromatin (**Figure 3 A**). Additional factors that drive early differentiation (Tcf4/7, Klf10, Rbpj, Pax2/8 and Uncx) and those that specify the proximo-distal lineage fates (e.g., Hoxb2/b4, Lhx1, Hnf1b, Pou3f3, and Klf6 and Elf1/5) are subsequently engaged. Nearing the end of their lifespan, P2-NPCs are actively involved in differentiation, and this is reflected by a TF signature mostly composed of key fate-determining factors of proximal and distal fates, such as Hnf1b, Hnf4a, Hoxd8 and Elf family members (**Figure 3 B**). Interestingly, Nfkb, recently identified by single cell analysis of human kidneys as a potential regulator of proximal tubular integrity (20), is represented (along with its partners Rel-a and Rel-b) in the trajectory (**Figure 3 B**).

**Figure 3.**
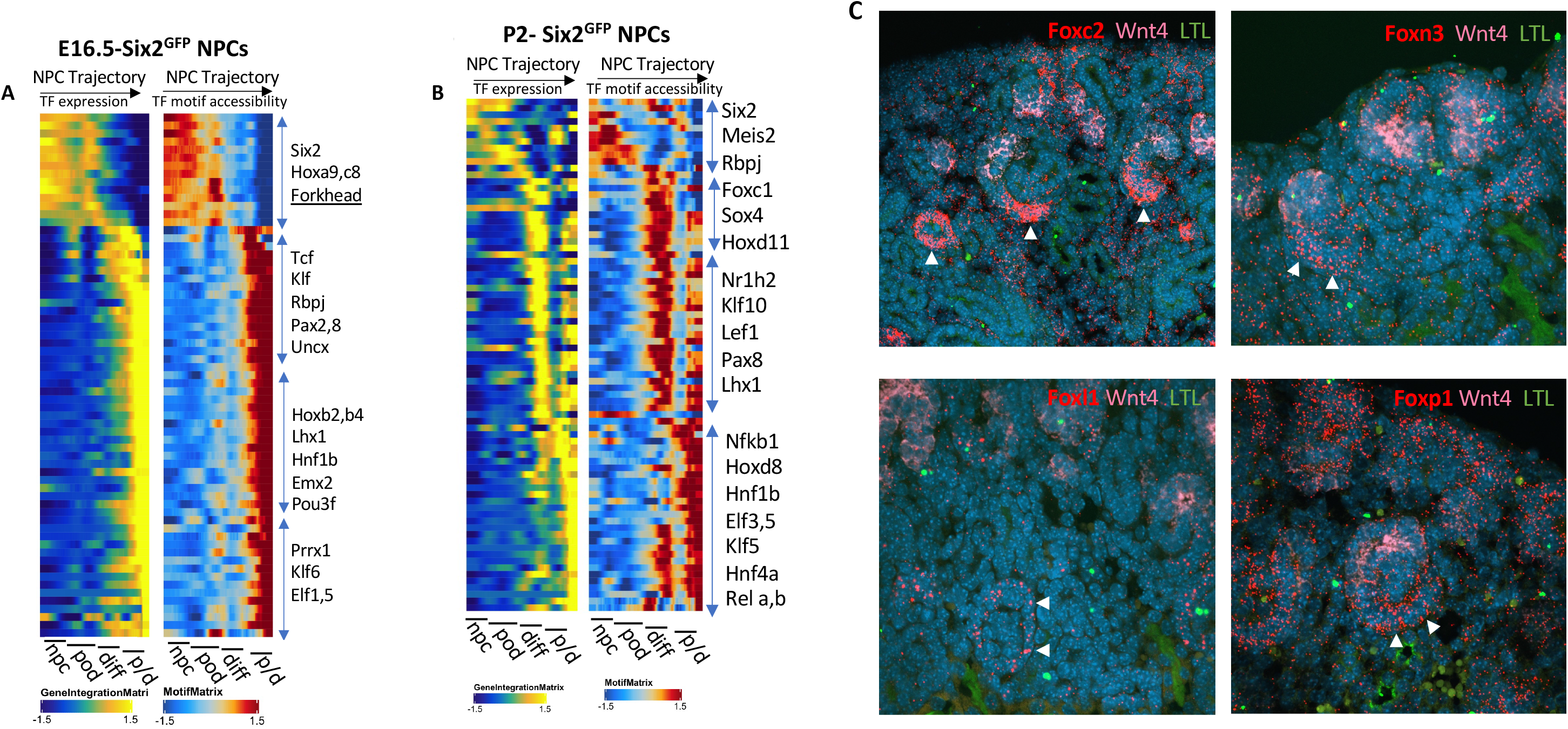
Six2^GFP^ cellular trajectory based on matched transcription factor expression and motif activity. (A,B) Side-by-side heatmaps of TF gene expression and ChromVar motif scores across pseudotime and cell states. (C) Confocal microscopy images (x60 magnification) of section RNA in situ hybridization (RNAscope) showing expression of Foxc2, *Foxl1*, *Foxn3* and *Foxp1* in podocytes in E16.5 kidney (white arrows).

The early engagement of Foxn3 and Foxp1 factors in the NPC lineage was intriguing and reminiscent of Foxl1, recently identified in a similar analysis done at P0 (11). So, we explored where these factors might be expressed in the developing kidney. High-sensitivity RNA in-situ hybridization (RNAScope) revealed that Foxn3 and Foxp1 are expressed in the podocytes (albeit not exclusively) (**Figure 3 C**), whereas Foxo1 was diffusely expressed in the nephrogenic zone (not shown). We also confirmed that Foxl1 is enriched in the podocyte, as recently reported (11) (**Figure 3 C**). These findings imply a potential role for this subset of Fox factors in the specification of the podocyte fate.

### Multiome Analysis of the NPC lineage

To validate the above inferences based on integrated singleomes, we generated joint scATAC and scRNA datasets from E16.5 and P2 mouse kidneys using the Chromium Single-Cell Multiome ATAC+GEX platform (10x Genomics). Filtering across both data modalities yielded 7774 and 10,765 cells with high quality transcriptome and epigenome profiles, respectively (**Figure S8**). We obtained 168,566 and 171,147 accessible peaks representing potential CREs at E16.5 and P2, respectively. We clustered the scATAC-seq and scRNA-seq datasets and annotated these clusters using gene expression and gene scores, then selected the cell clusters representing the nephrogenic (NPC) lineage (npc-to-distal) for further analysis. We next integrated the derived gene scores (chromatin accessibility) with gene expression scores (GEX) (**Figure 4 A-D**). Key nephrogenesis factors such as *Six2* and *Pax2* (NPC), *Foxc2* (podocyte), *Hnf4a* (proximal), *Pou3f3* (LOH), and *Gata3* (distal tubule) showed strong cluster specific enrichment using gene and GEX scores (**Figure 4 E**). Marker-based cell clustering of chromatin accessibility defined the cell types of the nephrogenic lineage (**Figure 4 F, Figure S9**). Visualization of accessible peaks across all clusters (C1 to C17) in the *Six2* and *Hnf1b* gene loci (+/-70 kb from the TSS) revealed enriched co-accessible CREs in NPCs and proximo-distal clusters, respectively (**Figure 4 G**). By comparing gene scores and GEX scores, we observed two prevalent patterns: type I genes were characterized by regulatory chromatin opening preceding gene activity. Examples in this group of genes are transcriptional regulators, enzymes, and G-protein coupled receptors (**Figure S10)**. In group II genes, regulatory chromatin opening coincides with gene activation. The latter group is enriched in channels and transporters. The biological significance of these differential chromatin behaviors is not clear but may be important in lineage priming and terminal differentiation.

**Figure 4.**
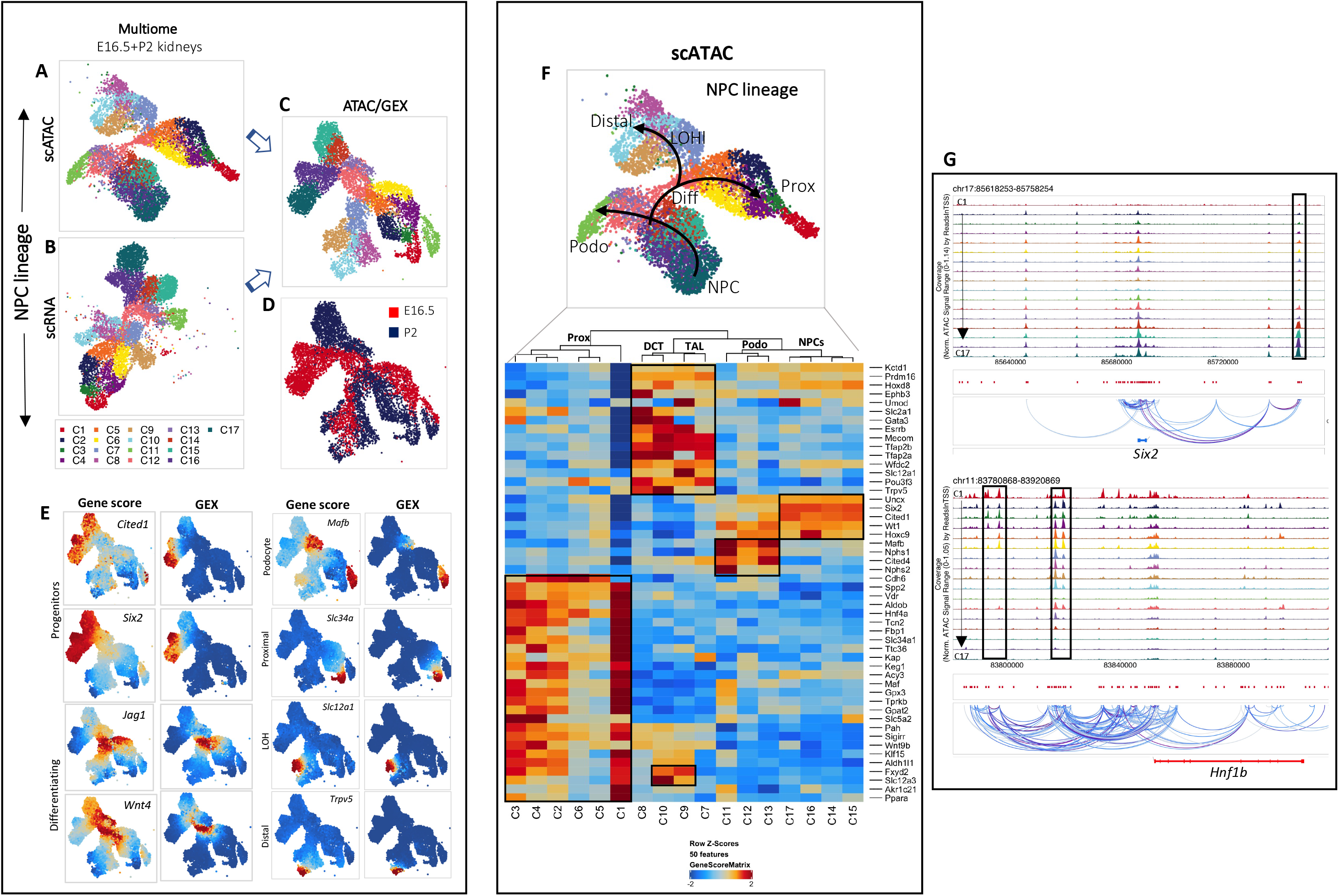
Multiome single-cell ATAC-seq/Gene expression (GEX) of E16.5/P2 kidneys. (A) scATAC UMAP-based clustering. (B) scRNA UMAP-based clustering. (C,D) Matched ATAC/RNA UMAP-based clustering. (E) Heatmap of gene scores and GEX of representative cell-type specific genes. (F) UMAP plot (top) and heatmap (bottom) representing E16.5/P2 chromatin accessibility-based assignment of cell clusters. (G) Representative chromatin accessibility tracks of all clusters for the *Six2* and *Hnf1b* genes. Boxed areas denote gene regulatory regions.

### Motif enrichment analysis reveals maturational differences in TF activity

We next performed ChromVAR motif enrichment analysis on marker peaks and differentially accessible peaks to determine if these groups of peaks are enriched for binding sites of specific TFs (Cutoff = FDR<=0.1 and Log2FC>0.5) (**Figure 5 A, B**). We found that bHLH motifs are enriched in podocyte and NPCs. In comparison, homeobox motifs are most enriched in NPCs, while Forkhead box motifs are enriched in podocytes. Distal clusters were enriched with accessible Pou motifs, whereas proximal clusters were enriched in Hnf4/Hnf1/Rxr motifs. AP-1 motifs were enriched in intermediate clusters 2, 5 and 8 representing differentiating proximal and distal tubules. Ranking the top differentially accessible motifs based on ChromVAR variability scores across all cells revealed Hnf1 motifs are the most represented in the accessible peaks (**Figure 5 C**). Other TF motifs that are also overrepresented include Six, Hnf4 and Hox motifs. Our previous bulk ATAC-seq analysis revealed that older (P0-P2) NPCs have greater chromatin accessibility to AP-1 factors. Here, we confirm through differential analysis of scATAC seq profiles that NPC maturation is associated with a remarkable shift in TF accessibility from the “progenitor” type (Six2, Hox, Wt1, Fox) to AP-1 factors (**Figure 5 D**). Other TFs that gain increased accessibility in P2 are terminal differentiation genes including Hnf1, Tead, Nfkb and Hnf4.

**Figure 5.**
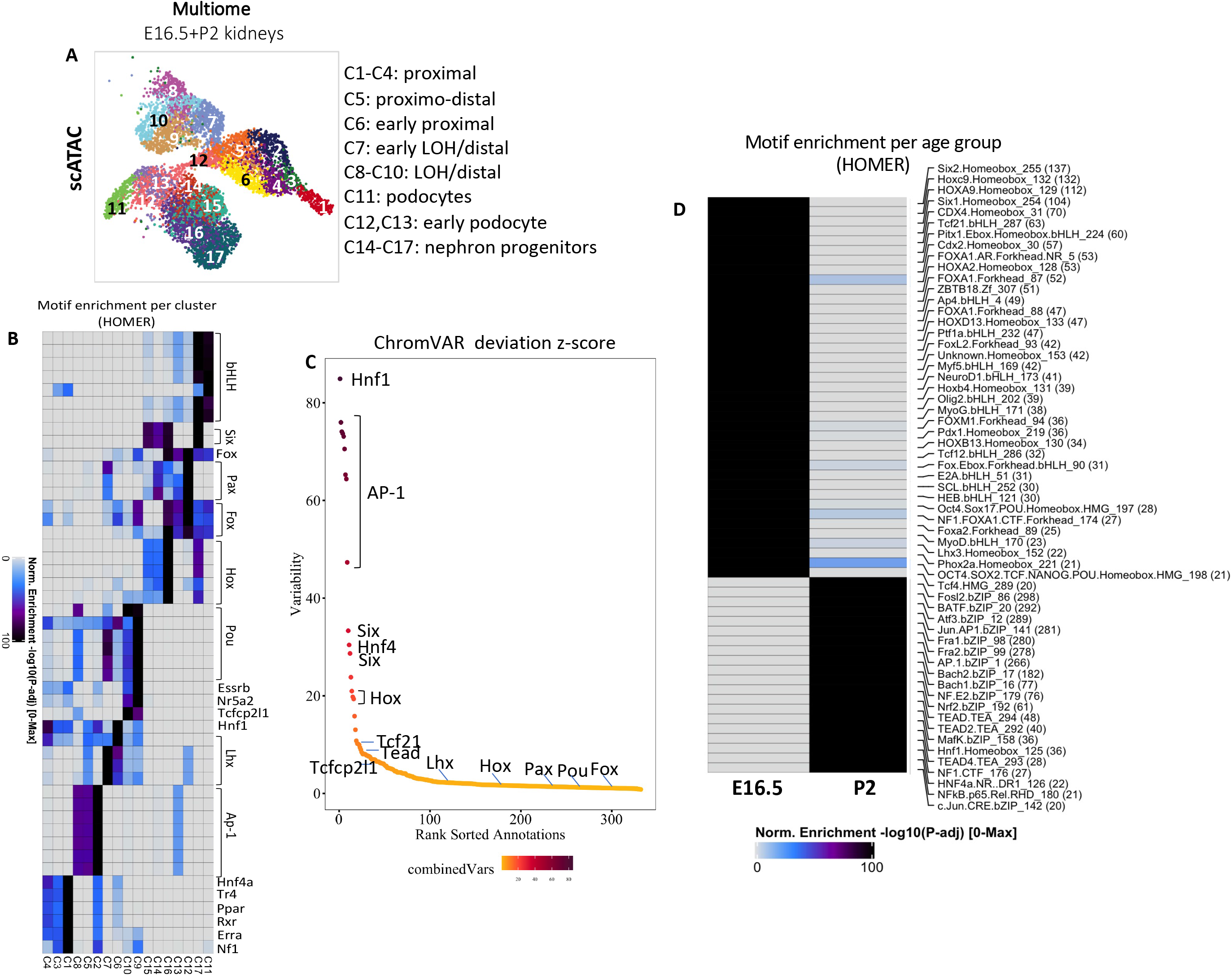
Multiome analysis of differential chromatin accessibility in E16.5/P2 NPC lineage. (A) scATAC-based UMAP plot with cell-type specific annotations. (B) ChromVAR TF motif enrichment per cell cluster. (C) ChromVAR deviation scores for the most represented TF motifs in NPCs. (D) Age-based motif enrichment analysis in E16.5 and P2 NPCs.

### Trajectory of TF motif activity

Integration of TF gene expression with CRE accessibility along the NPC differentiation trajectory illustrated the dynamic accessibility to fate-determining TFs in the progenitor state (Six2, Meis2/3, Tcf21, Hoxc8/a9/a11), committed (E2f6, Stat3, Sox12, Pax2, Pax8), podocytes (Wt1, Fox factors), proximo-distal committed cells (Nfkb1, Essrb, Elf3, Klf6, Hoxb2/b4, Lhx1), distal (Pou3f3), and proximal fate (Hnf4a/g, Hnf1b, Ppara, Nr1h4) (**Figure 6**). To gain further insights into cell type-specific regulons (groups of co-regulated genes), we examined TF regulon activity using <u>S</u>ingle-<u>C</u>ell r<u>E</u>gulatory <u>N</u>etwork Inference and <u>C</u>lustering (SCENIC). SCENIC incorporates TF cell type specific expression with motif-based filtration to infer potential direct targets of each TF (regulons). Regulon activity was quantified and was binarized into “on” or “off” based on activity distribution across cells. As shown in **Figure 7 A and B**, SCENIC clustered the cells based on the regulon states of each cell showing strong enrichment of *Meis1, Zeb1, Hoxa10, Hoxd11*, *Ezh2, Foxc1*, and *Tcf7l1* regulon activity in nephron progenitors, *Mafb* and *Lef1* in podocytes, *Emx2, Hnf1b, Pax8, Lef1, Sox9, Nfkb1* in differentiating cells, *Tfdp1, Brca1, E2f* factors in cycling NPCs, *Hnf1a, and Hnf4g* in proximal tubules, *Atf3 and Paprgc1a* in loop of Henle, and *Hoxd8, Gata3* and *Pou3f3* in distal nephron, respectively. Examples of regulon activity and corresponding TF gene expression are shown in **Figure 7 C**.

**Figure 6.**
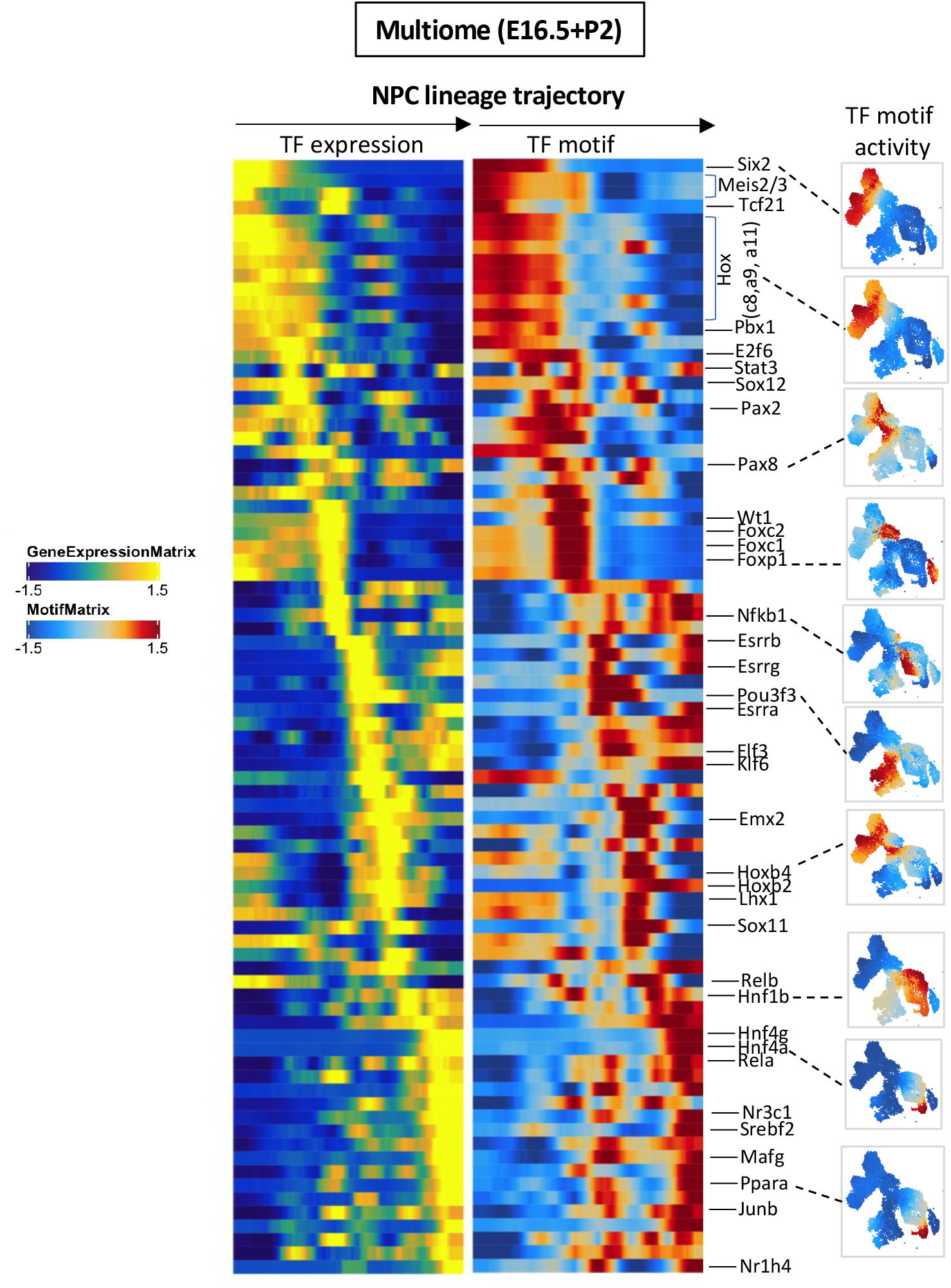
Multiome NPC trajectory based on matched transcription factor expression and motif activity. Side-by-side heatmaps of TF gene expression and motif scores across pseudotime.

**Figure 7.**
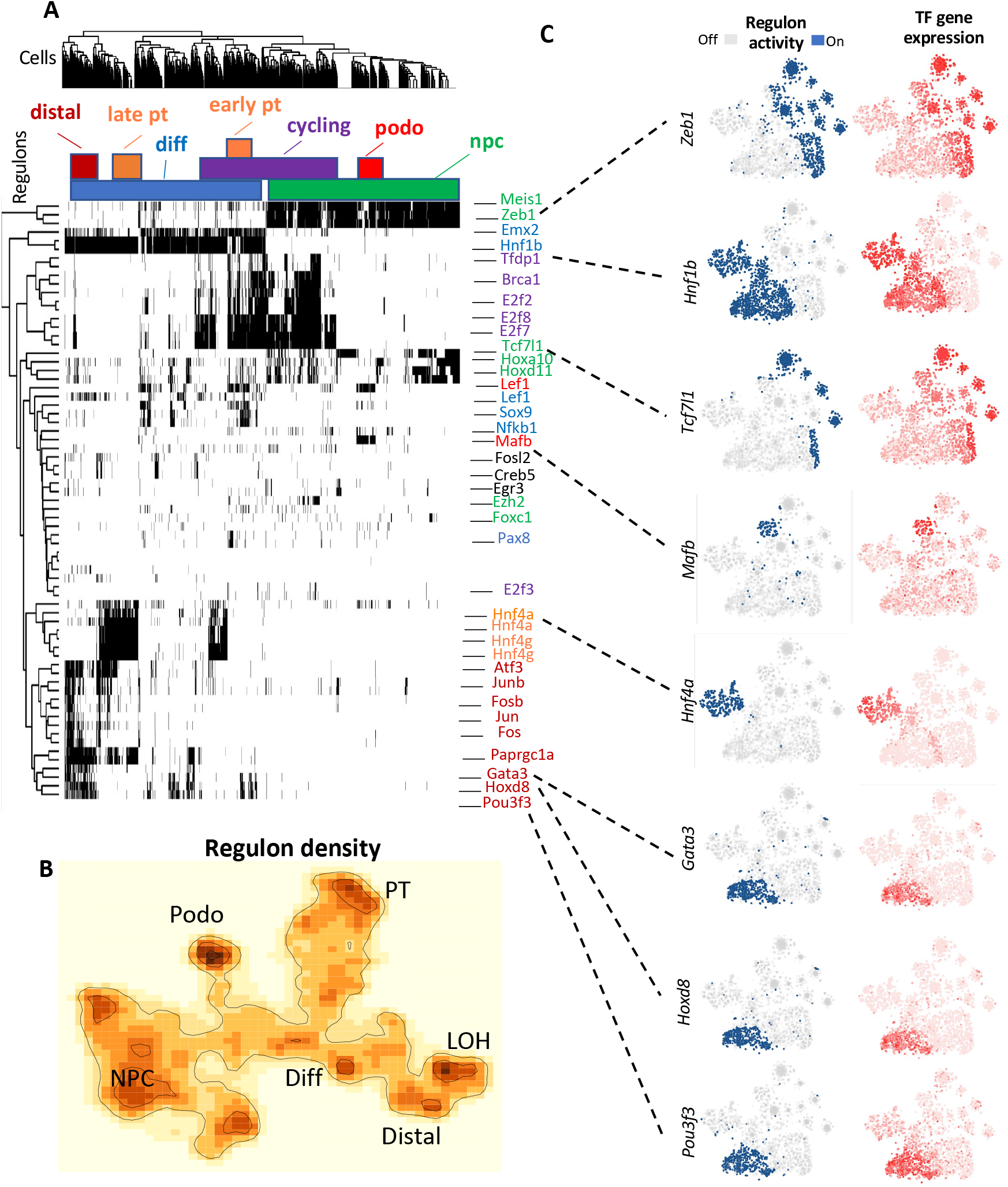
Regulon activity of the NPC lineage. (A) SCENIC-generated heatmap of cell type-specific regulons. Regulons are binarized to “on” (black) or “off” (white). The genes are colored by the cell cluster. (B) tSNE plot of regulon density representing regulon states as inferred by SCENIC algorithm. (C) tSNE plots showing regulon activity of selected regulons for nephron progenitors (*Zeb1, Tcf7l1*), differentiating (*Hnf1b*), podocyte (*Mafb*), proximal (*Hnf4a*), and distal (*Gata3, Hoxd8, Pou3f3*) tubules.

### CRE-gene associations across the NPC lineage uncover specialized cellular functions

Like our analysis in Six2^GFP^ cells (Figure 2), we applied a correlation-based approach that links CREs to gene expression, identifying 24270 dynamic interactions across all E16.5 and P2 NPC lineage cells and grouped these interactions into 10 k-means clusters (**Figure 8 A**). GREAT analysis of linked genes clustered well into various cell functional types such as metabolic processes, segmental nephron morphogenesis, glomerular/podocyte differentiation and tubular transport (linked genes are listed in Supplemental tables 1-10). Of significant interest, CRE-gene linkages identified genes involved in highly specialized cell functions such as macula densa, mesangial and podocyte morphogenesis.

**Figure 8.**
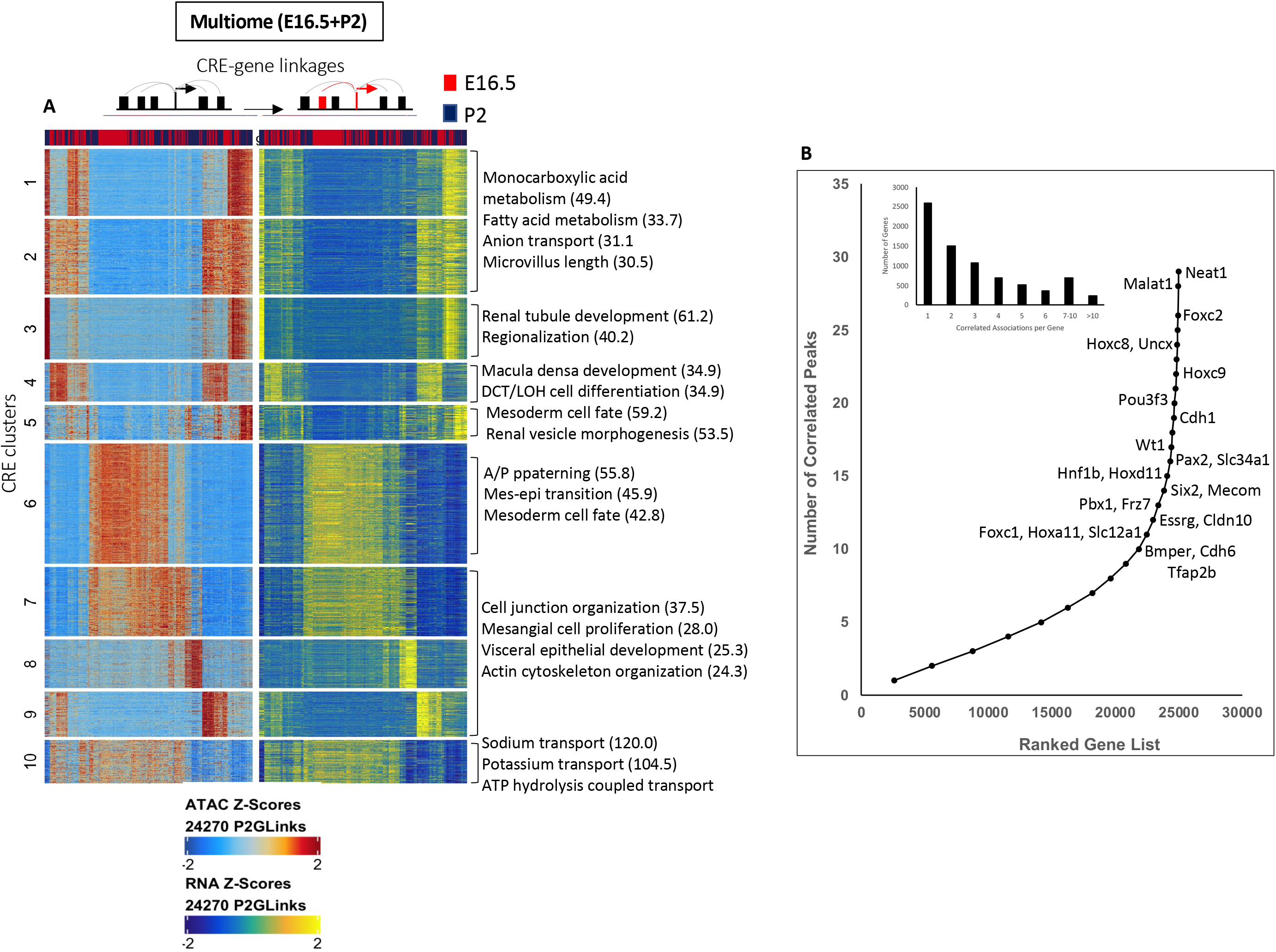
Multiome analysis: inferring regulatory chromatin and TF trajectories. (A) Correlations of 24270 CRE-gene associations ATAC and RNA Z-scores across all cell states (k means=10). rGREAT annotation of cluster-related biological processes. (B) Inset, bar graph depicting ranking of genes based on the number of correlated peak-gene associations (+/-250 kb from TSS). Line graph depicts ranking of genes harboring chromatin domains with a minimum of 10 peak-gene links.

Using a minimum of 10 CRE-gene links per gene, we defined a set of key developmental TFs and genes that define NPC, podocyte, committed, proximal and distal fates such as *Wt1, Hox, Six2, Foxc2, Hnf1b, Lef1*, *Cdh6, Mecom, Tfap2b,* and *Pou3f3* (**Figure 8 B**), illustrating the value of CRE-gene associations in identifying lineage- and fate-specific genes, and validating the results obtained in Six2^GFP^ cells. Interestingly, the top 2 ranked genes with the highest number of CRE-gene links are *Neat1* and *Malat1*; both genes encode long non-coding RNAs postulated to function in renal epithelial cell protection and repair pathways (21).

### Analysis of early NPCs recovers Fox gene activity

To further understand the chromatin landscape of NPCs, we sub-clustered the Cited1-enriched cell population in E16.5 NPCs and obtained 4 cell clusters (**Figure S11 A**). UMAP plots of gene scores and gene expression revealed that C2 is enriched in *Cited1* and *Six2* (**Figure S11 B**). Also, Six2 motif activity is high in C2. In comparison, C4 is enriched in *Fox* family members (*Foxc2, Foxl1, Foxp1 and Foxn3*) when assessed by gene scores, gene expression and motif activity (**Figure S11 B**, arrows). Interestingly, Wt1 motif activity is also high in C4. Examination of H3K27ac ChIP-seq and ATAC-seq tracks (GSE124804) revealed that Six2 and WT1 bind the *Foxl1* gene at accessible chromatin regions (peaks) upstream of the TSS (**Figure S11 C)**, suggesting that Six2 and Wt1 may directly regulate the cell type-specific expression of Fox factors. Analysis of TF motif enrichment in accessible peaks showed that C2 is enriched with accessible motifs for Six2, and the AP-1 family, while C4 is enriched in Fox motifs (**Figure S11 B, D**). Furthermore, matching TF expression with motif activity across pseudotime revealed high AP-1 and p53 activity (cell cycle/survival) in the beginning of the trajectory as compared to high Fox, Wt1 and Klf15 (podocyte) activity at the end of the trajectory (**Figure S11 E**).

In summary, elucidation of the accessible chromatin landscape in nephron progenitors informs the mechanisms of NPC choices to alternate fates and provides a foundation for future studies in disease states characterized by abnormal nephrogenesis.

## Author Contributions

**S.H.** analyzed the scATAC and multiome sets and participated in manuscript writing and figure preparation; **G.T.** analyzed the scRNA data and assisted in scATAC analysis and figure preparation; **H.L.** participated in kidney cell processing and FACS analysis; **C-H.C.** participated in mouse breeding, genotyping, tissue preparation, and conducted the in situ hybridization studies; **S.E-D.** designed the experiments and participated in writing the manuscript and preparation of figures.

## Acknowledgments

Special thanks to the members of the Tulane Center for Translational Research in Infection and Immunity for single-cell sequencing and for the Confocal Microscopy facility, and the Humphrey laboratory for providing the computational algorithms for plotting CCANs using Circlize visualization in R (0.4.11).

## Data Sharing

All data related to scATAC-seq, scRNA and multiome datasets reported in this paper have been deposited to the Gene Expression Omnibus (GEO) under accession number GSE180902 and GSE124804.

---

**Supplemental Figure 1.**
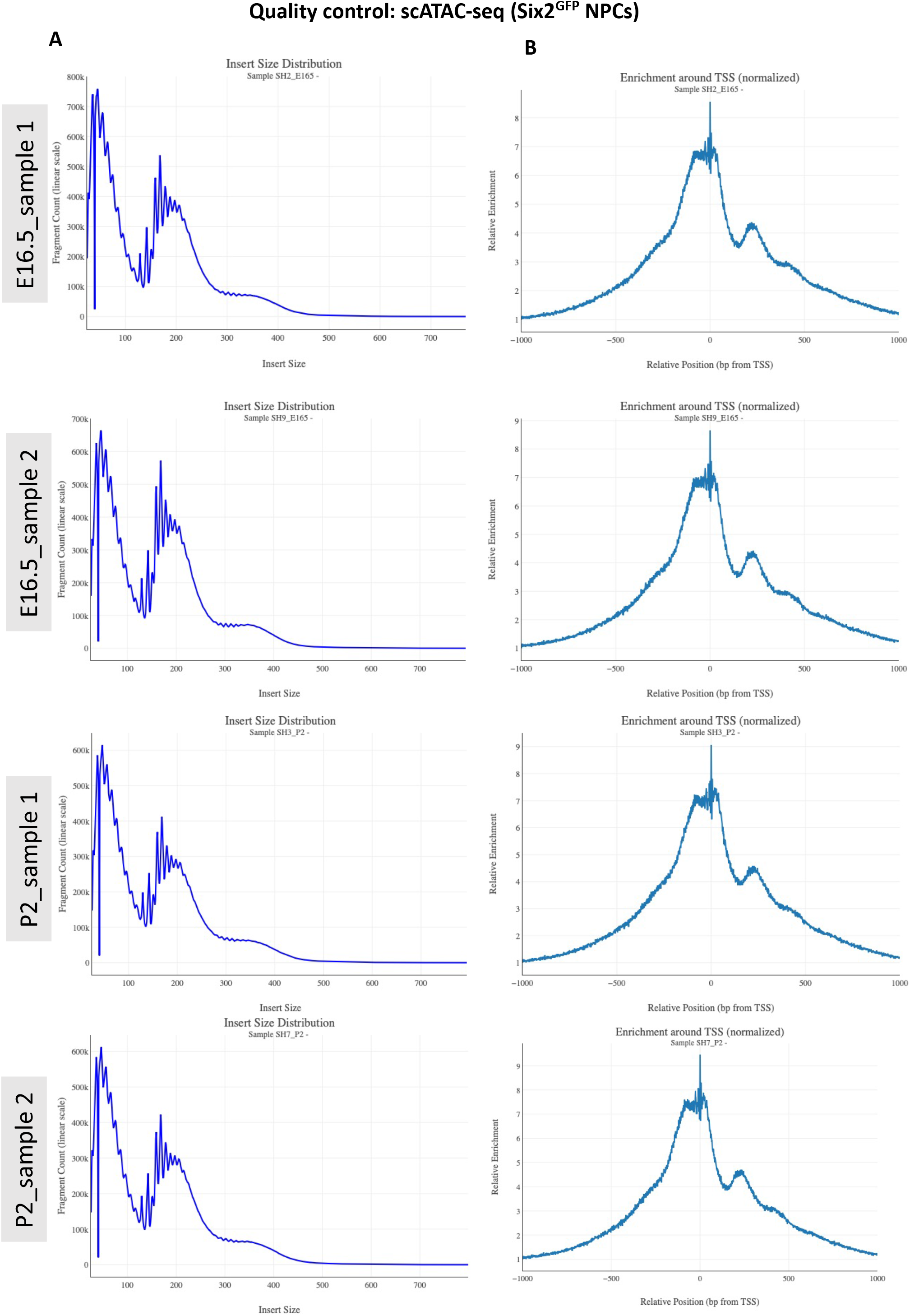
Quality control for the scATAC-seq data. (A) Insert size distribution of the scATAC-seq samples showing characteristic nucleosomal periodic patterns. (B) Transcription start sites (TSS) signal enrichment of the scATAC-seq samples.

**Supplemental Figure 2.**
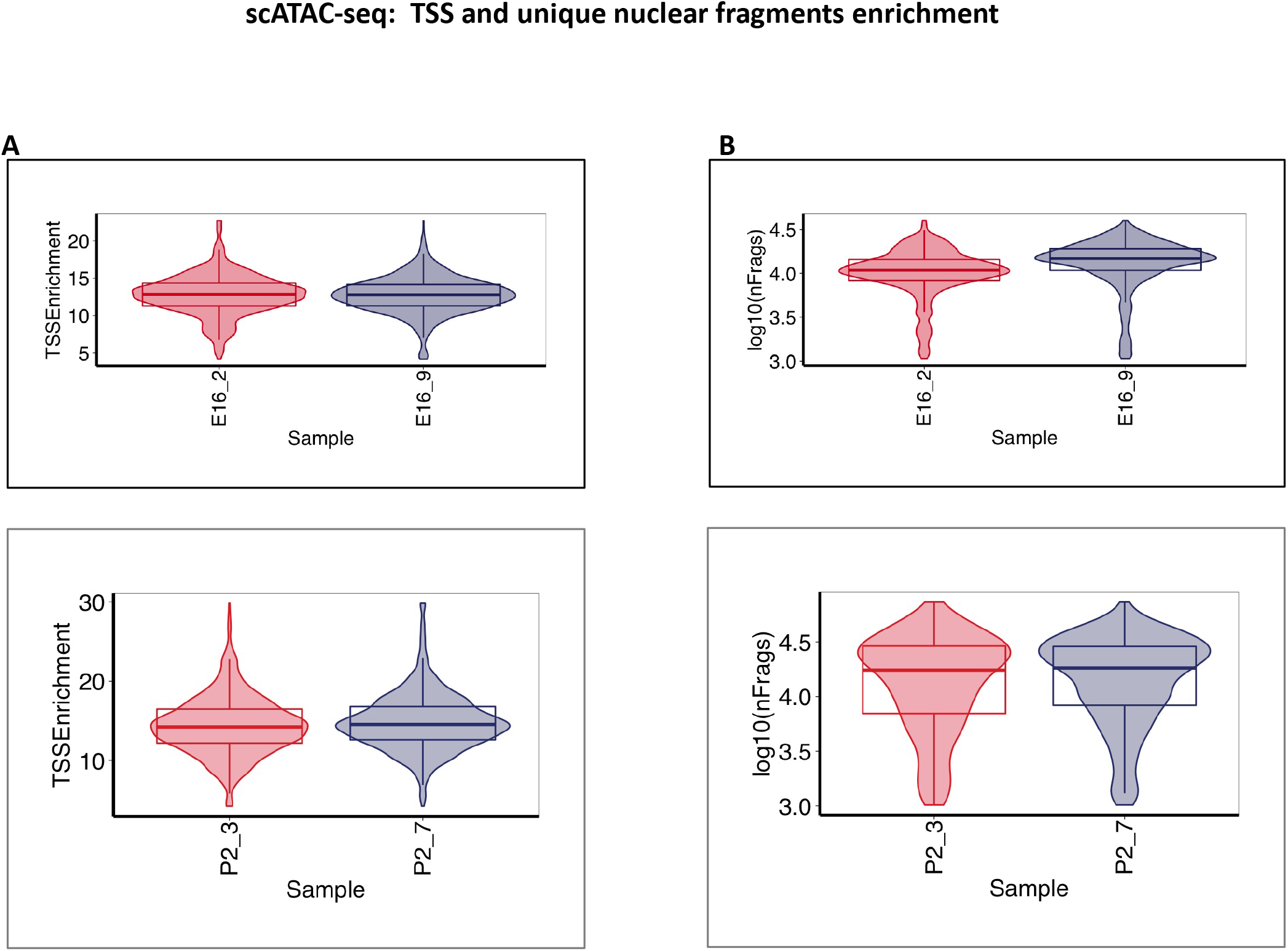
TSS and unique nuclear fragments enrichment in scATAC-seq data. (A) Violin plots showing TSS enrichment scores per sample at E16.5 and P2. (B) Violin plot showing the log(base10) of unique nuclear fragments per sample.

**Supplemental Figure 3.**
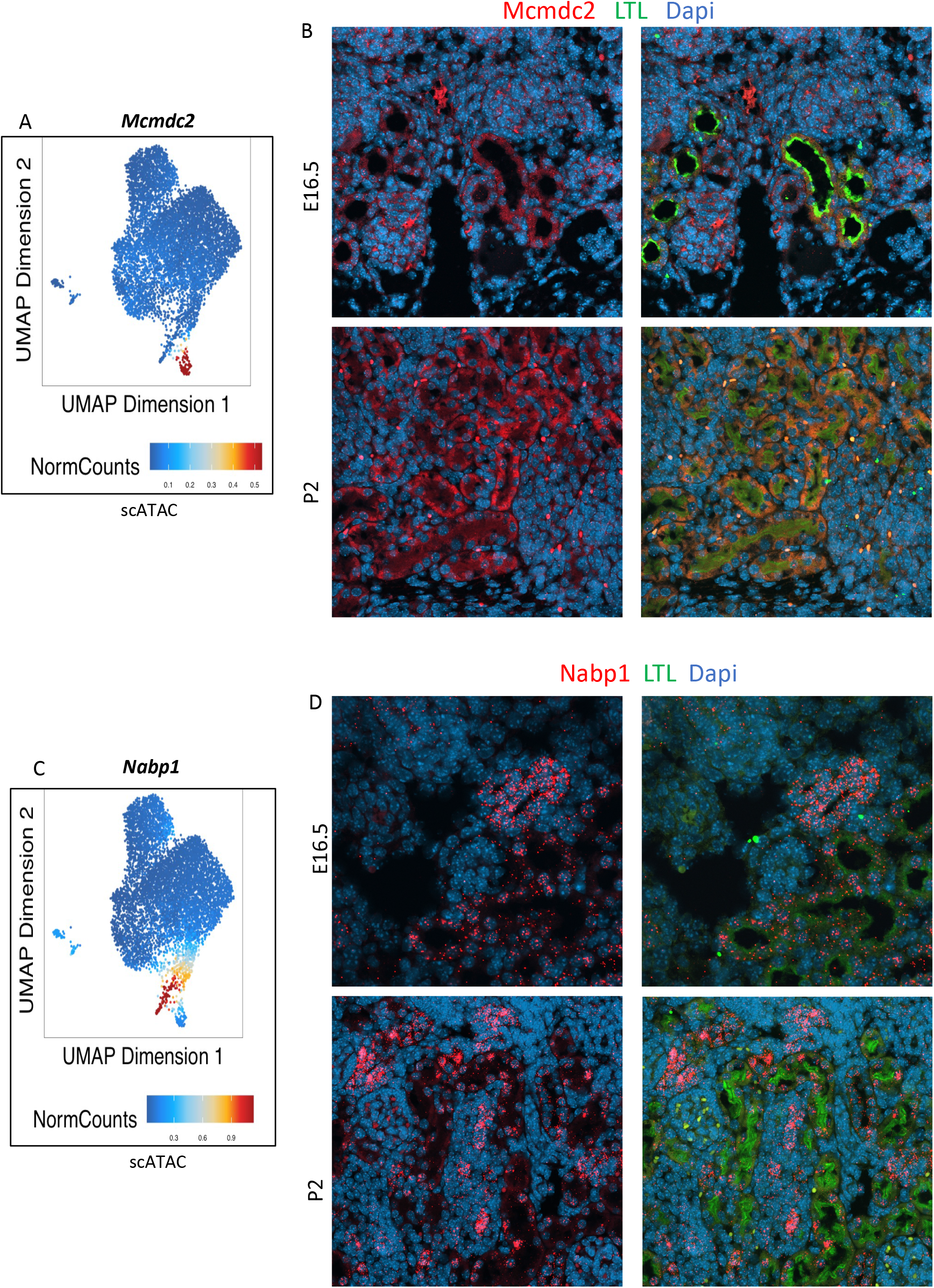
Chromatin accessibility gene scores identify new markers of proximal and distal tubules. (A,C) UMAP plots of P2 Six2^GFP^ scATAC depicting selective enrichment of chromatin activity for *Mcmdc2* and *Nabp1* in proximal and distal progenitors, respectively. (B) Confocal microscopy images (x40) of section in situ hybridization for *Mcmdc2* RNA in E16.5 and P2 kidneys showing predominant expression in proximal (LTL^+^) tubules. (D) Confocal microscopy images (x40) of section in situ hybridization for *Nabp1* RNA showing predominant expression in LTA-negative (distal) segments.

**Supplemental Figure 4.**
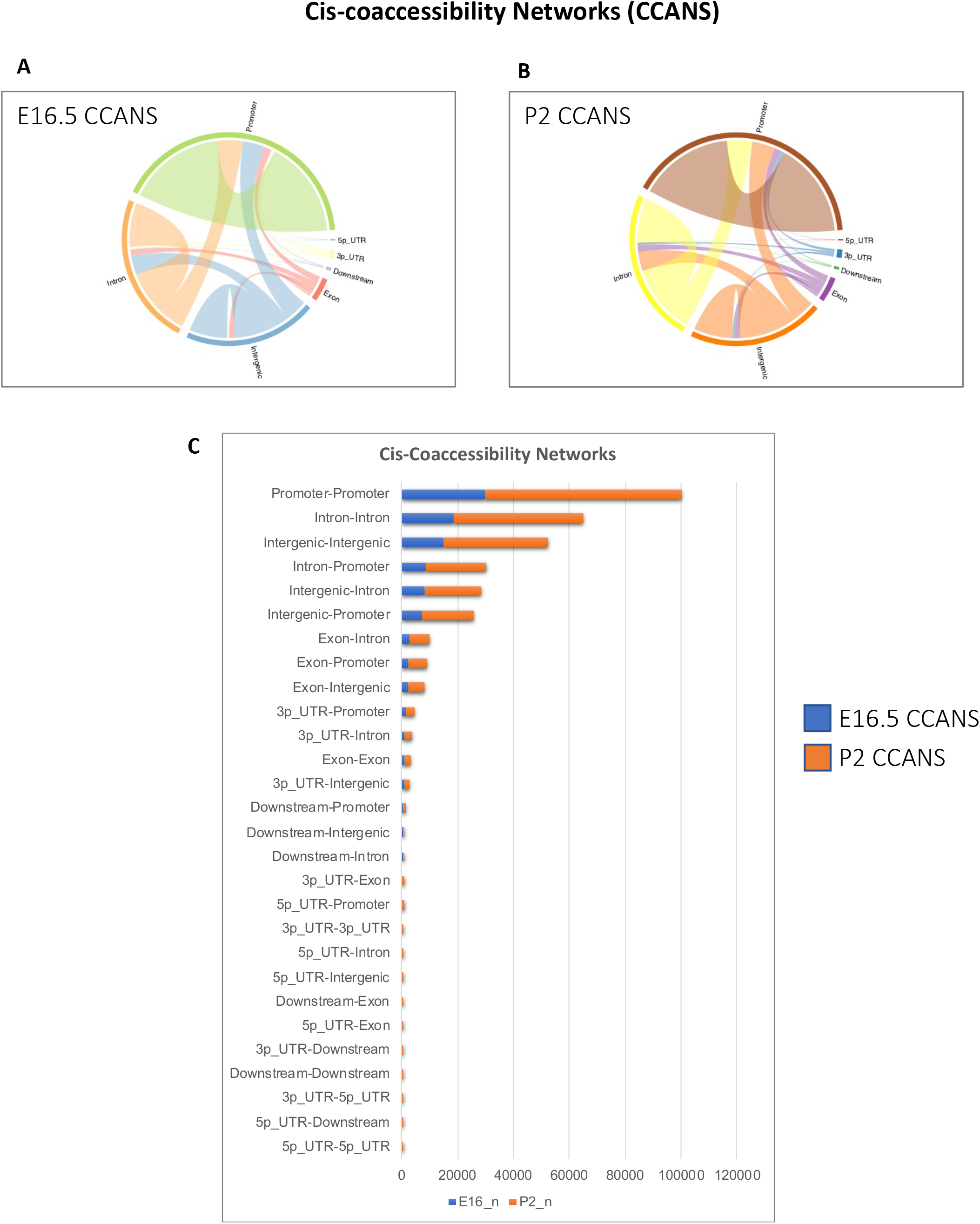
Cicero cis co-accessibility networks. (A,B) E16.5 and P2 Six2^GFP^ NPCs. Promoter-promoter, Intron-intron, Intergenic-intergenic connections are predominant in both age groups. (C) Age-related comparison of Cicero cis co-accessibility networks (CCANs) showing enhanced Cicero connections with age.

**Supplemental Figure 5.**
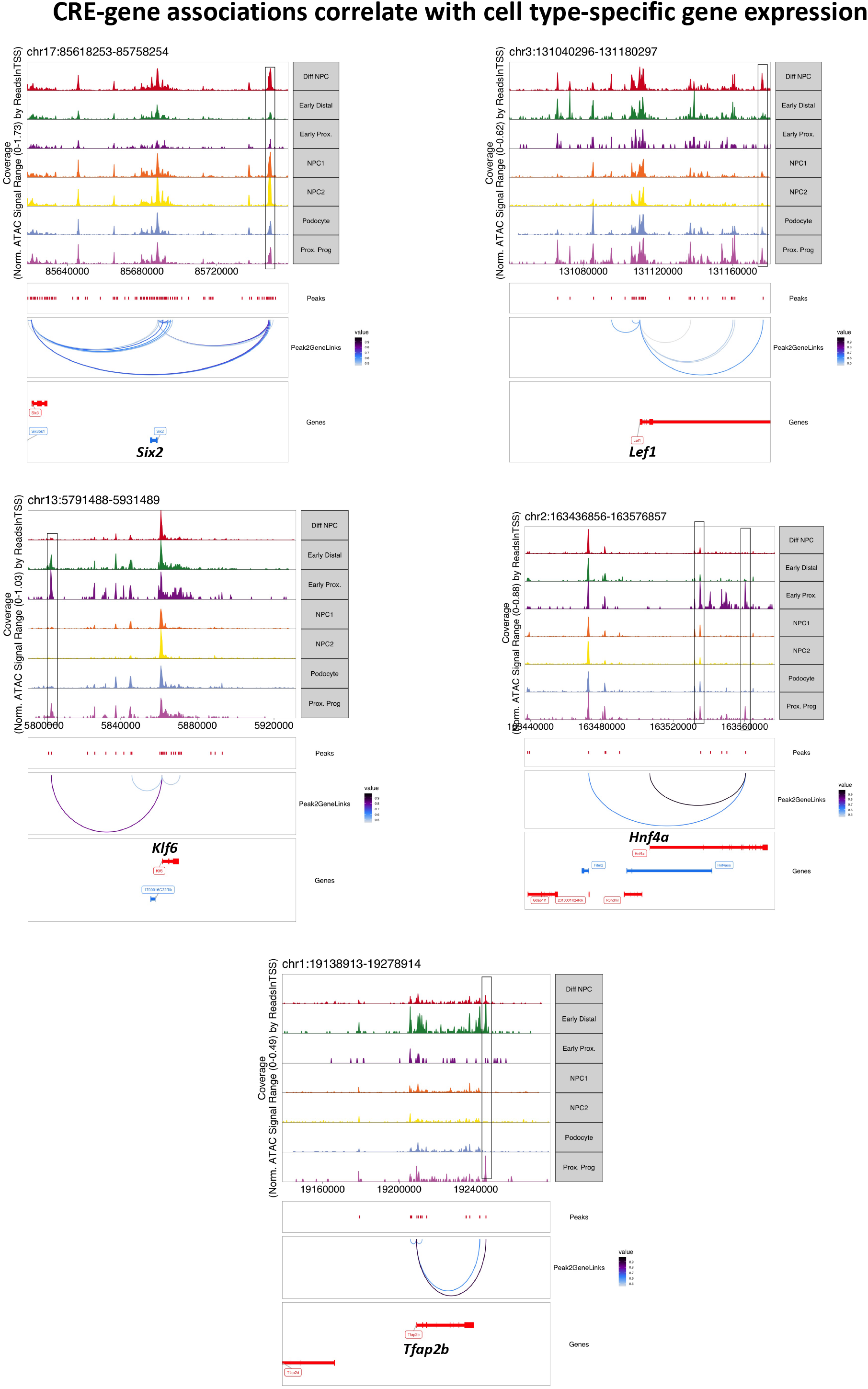
Peak-gene links representing co-variation in regulatory elements accessibility with gene expression and cell type-specific expression. Boxed areas depict differential peaks with significant correlation to gene expression. The strength of correlation is depicted by the color of the line.

**Supplemental Figure 6.**
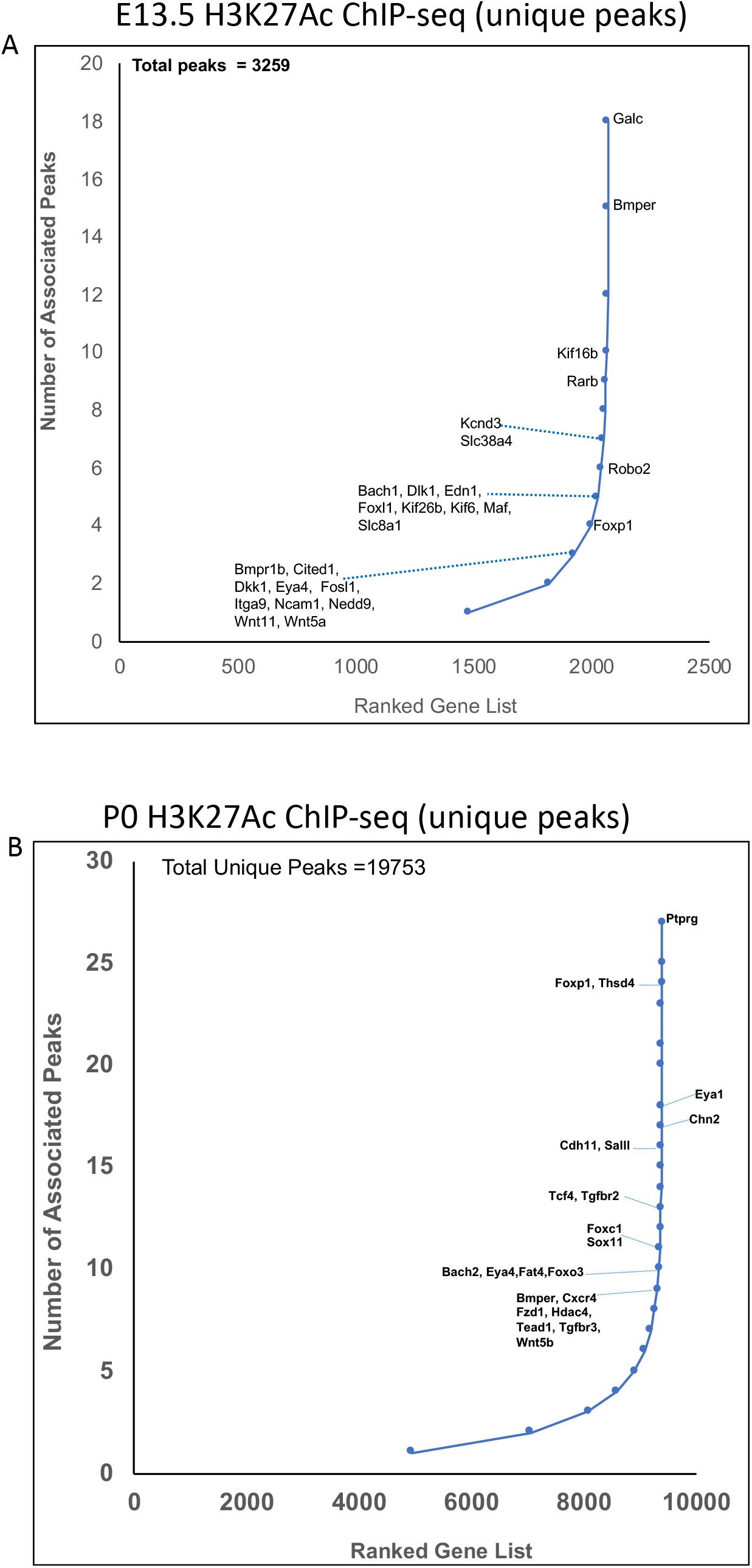
Ranking of ChIP-seq H3K27Ac-enriched peaks per gene in E13.5 and P0 NPCs. Datasets derived from (9)

**Supplemental Figure 7.**
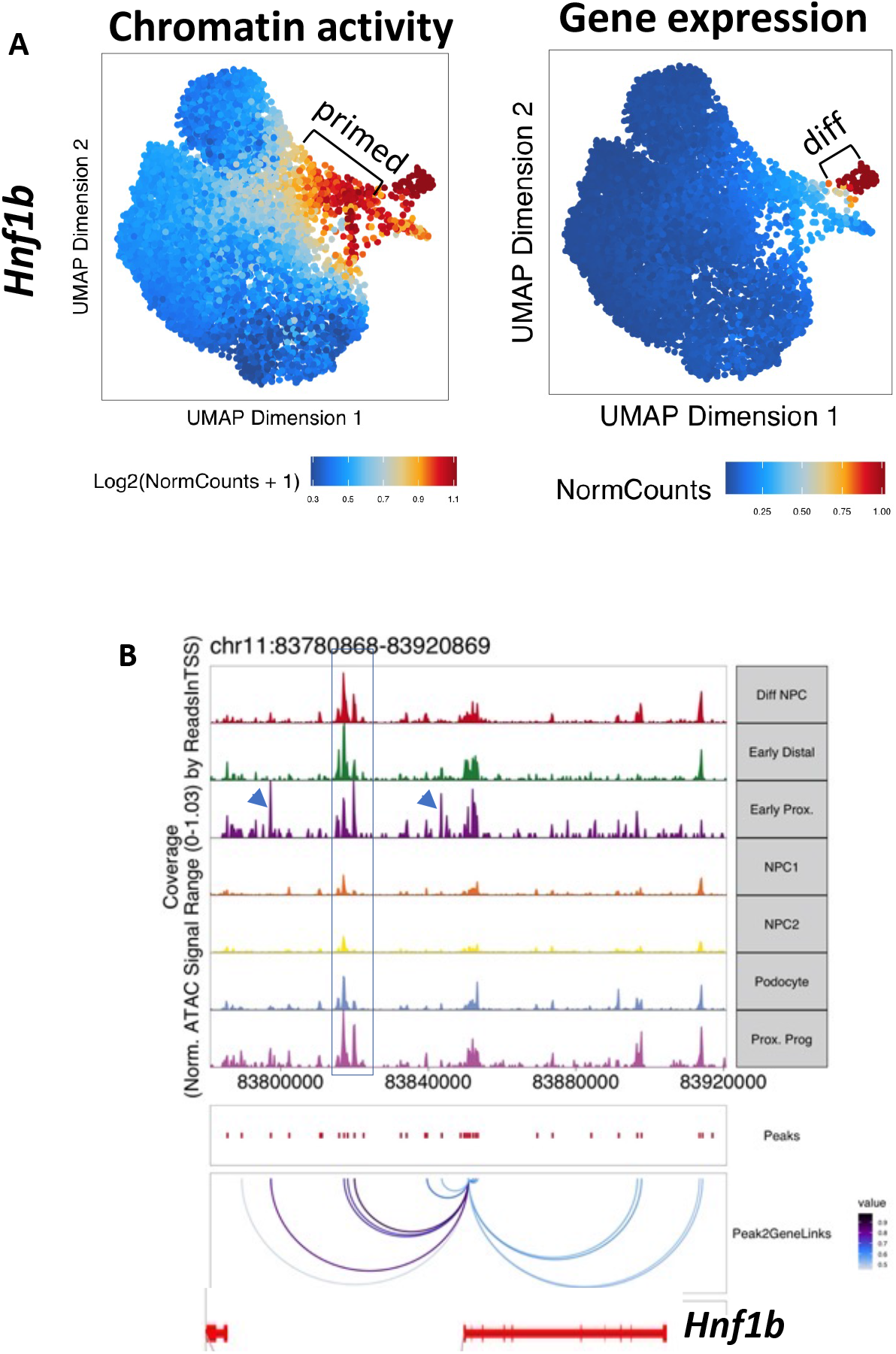
Chromatin activity and gene expression at the *Hnf1b* locus in P2 Six2^GFP^ NPCs. (A) The peak counts of all *Hnf1b* correlated peaks (left) and *Hnf1b* gene expression (right) colored in UMAP. The bracketed areas point to regions with differential signals. (B) Browser view of cell state specific peak-gene links at the *Hnf1b* locus. Putative enhancer peak-gene links are active in differentiating progenitors and those fated to become proximal/distal progenitors (box). Proximal-specific activity is associated with a gain of two additional peak-gene links (arrowheads).

**Supplemental Figure 8.**
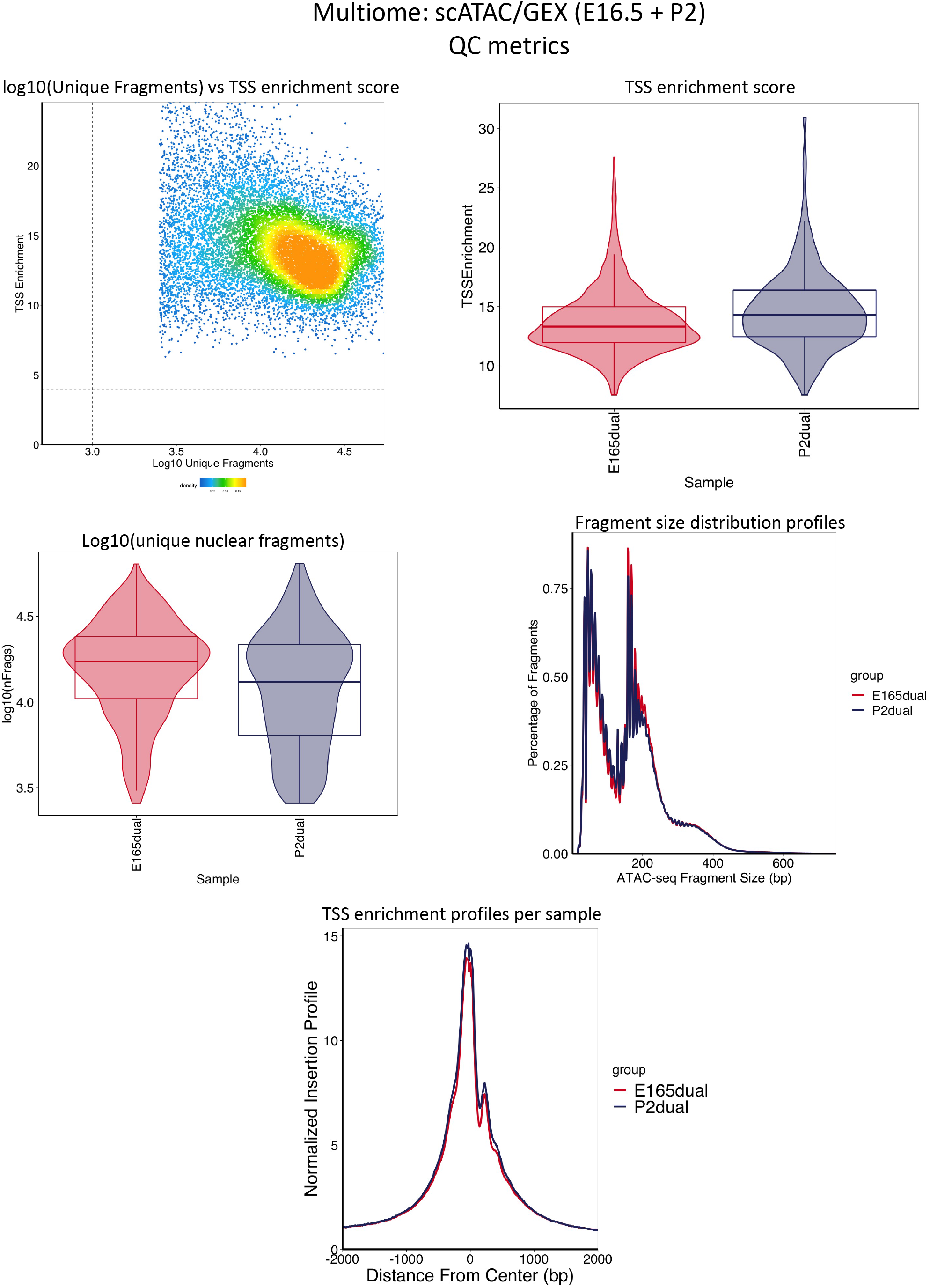
Quality control of Multiome scATAC/GEX (10x Genomics).

**Supplemental Figure 9.**
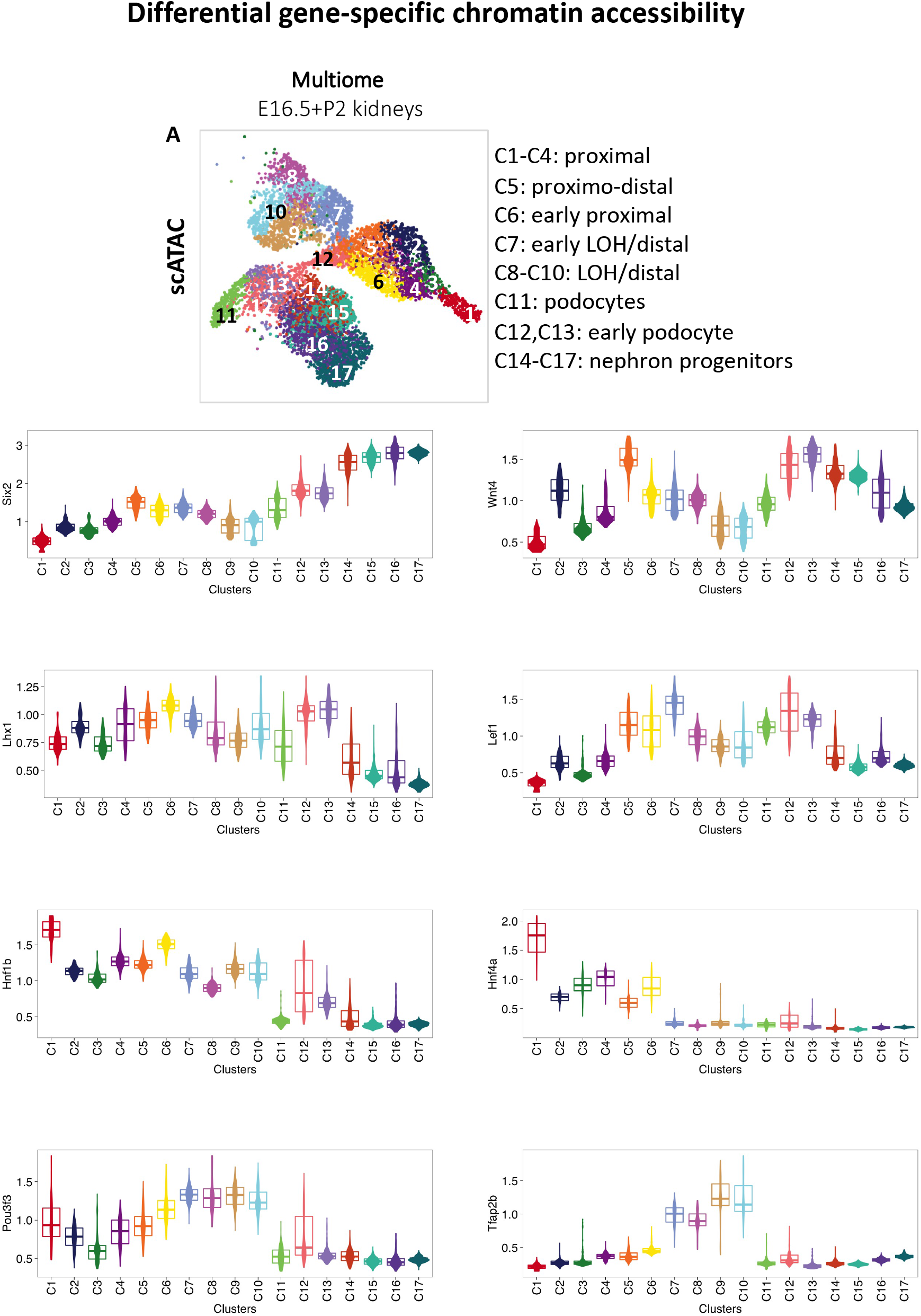
Differential gene-specific chromatin accessibility across cell clusters. Representative genes of progenitors (*Six2*), early (*Wnt4, Lef1*) and late (*Lhx1, Hnf1b*) differentiating, proximal (*Hnf4a*), LOH/distal (*Pou3f3 and Tfap2b*) segments are shown.

**Supplemental Figure 10.**
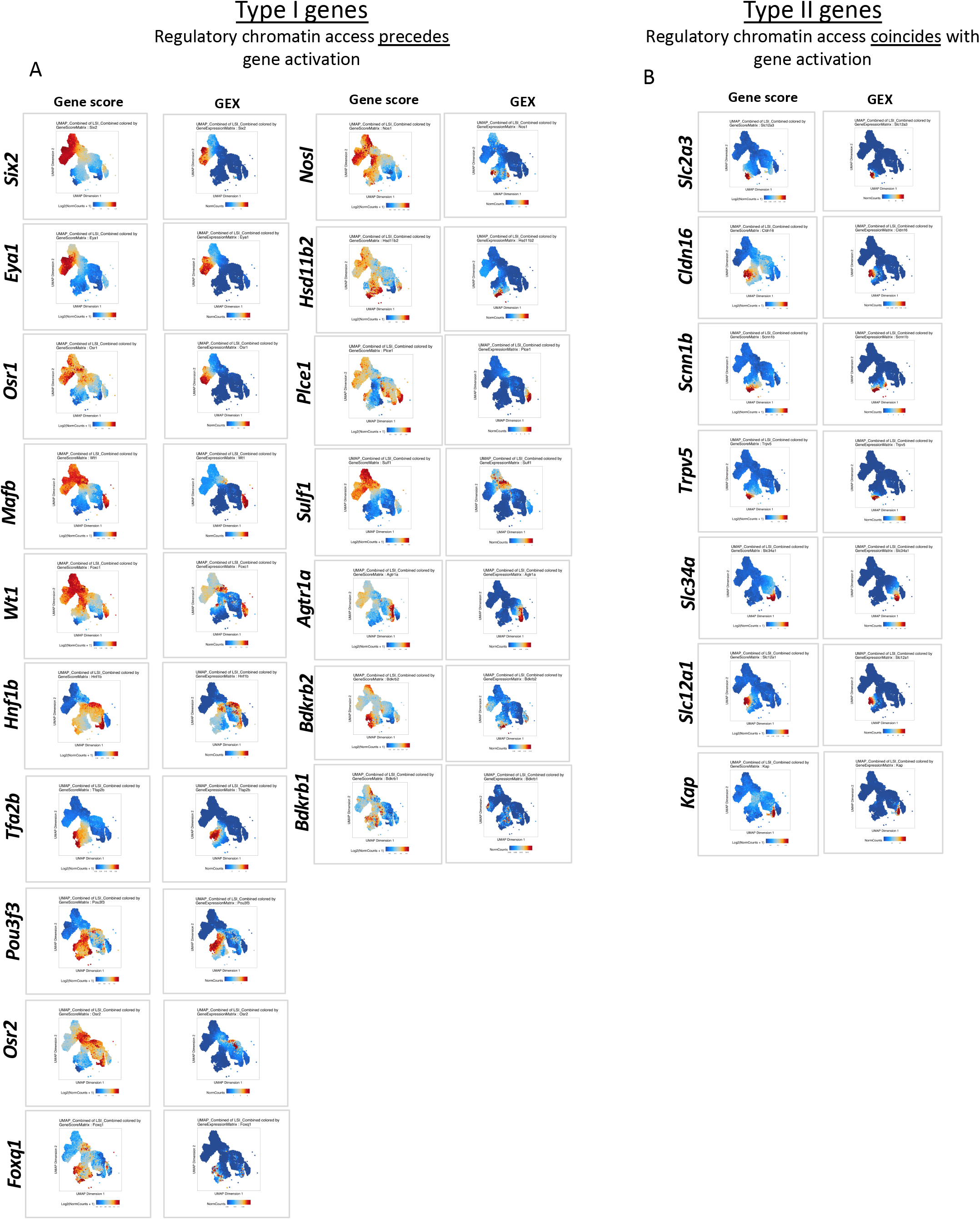
Multiome analysis of chromatin accessibility and gene expression defines two types of genes. (A) Type I: regulatory chromatin opening precedes gene activation. (B) Type II: regulatory chromatin opening coincides with gene activation.

**Supplemental Figure 11.**
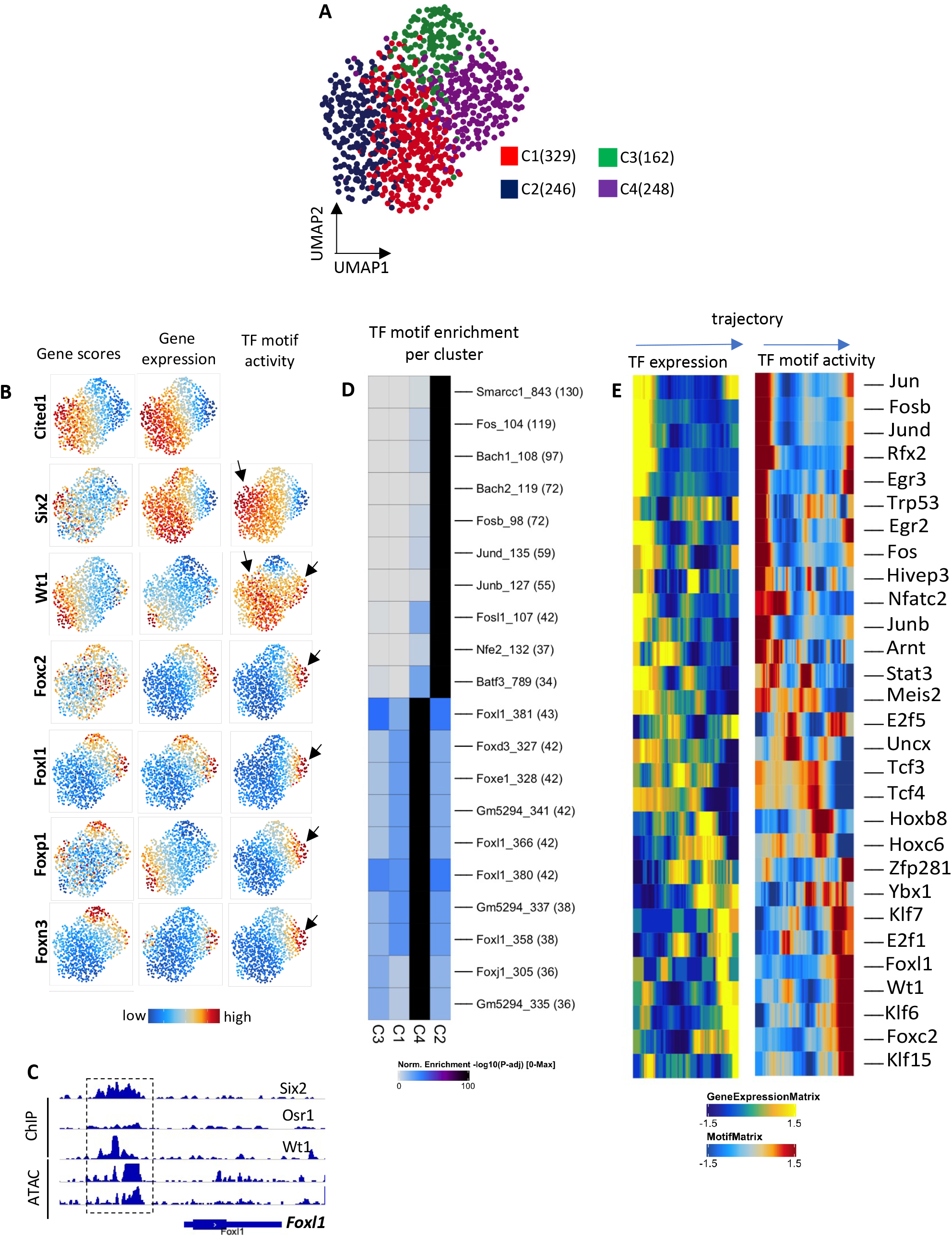
Chromatin landscape of the early progenitor cluster. (A) UMAP sub-clustering of E16.5 multiome Cited1-high cluster. Numbers in parentheses represent the cell number per sub-cluster. (B) UMAP plots highlighting chromatin accessibility (gene scores), gene expression and TF motif activity. *Fox* factors and *Six2/Cited1* have reciprocal cellular activity in C4 and C2, respectively. *WT1* motif activity overlaps with that of *Fox* factors. (C) Browser tracks of ChIP-seq (*Six2, Osr1*, and *WT1*) and bulk ATAC-seq in NPC-P0 (GSE804) highlighting the *Foxl1* locus. Six2 and WT1 co-occupy accessible chromatin regions upstream of the TSS (boxed area). (D) Enrichment of accessible AP-1 motifs in cluster 2 and Fox factors motifs in cluster 4. Note that the TF identity represents consensus elements and thus do not precisely reflect the exact member of the TF family. (E) Trajectory heatmap of TF expression and motif activity. Start of trajectory is cluster 2 and end is cluster 4.

